# Convergent gene family evolution underpins repeated transitions to metamorphic development across Pancrustacea

**DOI:** 10.64898/2026.05.06.723392

**Authors:** Giulia Campli, Ariel D. Chipman, Marc Robinson-Rechavi, Robert M. Waterhouse

## Abstract

Arthropod developmental modes are highly diverse, ranging from direct development with little morphological change between moults to metamorphic life-stage progressions characterised by profound transformations. Metamorphosis can be defined as a post-embryonic life-stage progression event leading to adulthood that is characterised by major morphological changes and modifications of the adaptive landscape. Within this framework, we compare four independent evolutionary life history transitions to metamorphic development across Pancrustacea. Using a phylogenomic dataset of 54 species spanning 26 orders, we investigated gene family evolutionary dynamics associated with the inferred origins of metamorphosis in Insecta, Copepoda, Eucarida, and Thecostraca. Compared with non-metamorphic sister lineages as well as descendent and ancestral nodes, transitions to metamorphic development were consistently associated with elevated gene family births and expansions. Although these expansions predominantly involved different gene families in each lineage, they repeatedly converged on shared biological functions, particularly those related to embryonic and post-embryonic development, morphogenesis, nervous system differentiation, and other processes relevant to the biology and evolution of metamorphosis. Evolutionary modelling further identified a subset of gene families exhibiting adaptive, lineage-specific expansions, including genes implicated in neural and sensory development, segmentation, and moulting. Together, these findings support a model in which independent transitions to metamorphic development repeatedly recruited different components of a shared developmental toolkit, achieving functional convergence through distinct genetic trajectories. This reframes the arthropod moulting programme as an evolutionarily flexible developmental substrate whose repeated modification has facilitated the emergence of complex multi-phasic life histories and contributed to the extraordinary diversification of Pancrustacea.

## Introduction

Life cycles comprising morphologically distinct developmental phases are widespread amongst animals. They occur in the majority of phyla, have evolved independently many times, and have persisted over vast evolutionary timescales (Moran 1994; Ten Brink et al. 2019). This phenomenon is evident within the clade Pancrustacea, comprising the subphylum Hexapoda and the paraphyletic crustaceans. The diversity of the clade is unparalleled both in terms of the number of described species, with more than one million from the class Insecta, and in terms of morphological diversity, with Crustacea including ten taxonomic classes and a remarkable diversity of body plans (Martin et al. 2014; Bernot et al. 2023; Chipman 2025). Pancrustacean lifelong developmental modes are highly diverse and span a range of two extremes: from minimal variation, such as water fleas with miniature hatchlings exhibiting no or little morphological differences between moults apart from growth in size, to maximal variation, where an extreme metamorphic transformation occurs as in butterflies (Bishop et al. 2006; Martin et al. 2014; Haug 2020a). Despite this diversity and lineage-specific morphological features that complicate comparisons across clades, there is a shared conceptual framework of metamorphosis: a post-embryonic life-stage progression that leads to adulthood, characterised by major morphological changes and modifications of the adaptive landscape (Bishop et al. 2006; Martin et al. 2014).

In Pancrustacea, metamorphosis is universally mediated through the process of periodic exoskeleton renewal, called moulting. The rebuilding of the exoskeleton allows not only the tissues within to grow in size, but also the formation of adult morphological traits. In Insecta, metamorphosis likely appeared in the last common ancestor of Pterygota, the winged insects (Jindra 2019; Truman 2019). In hemimetabolous insects, wingless immature stages possess a juvenile-specific cuticle type and have epithelial cuticle-secreting cells that are lost with adult wing maturation (Truman 2019). There are also striking distinctions in lifestyles between instars, for example, in the early diverging lineages of Ephemeroptera and Odonata, immature stages are aquatic, whereas adults are aerial (Truman 2019; Almudi et al. 2020). In fact, evolution of metamorphosis in insects likely represents a developmental strategy to accommodate the evolution of flight (Truman 2019). In holometabolous insects, wings develop at the end of the larval phase, from a special set of cells grouped in the imaginal discs. In hemimetabolous insects, a modification of post-embryonic development is also observed, as maturation of wing pads into articulated and functional wings is delayed to adulthood (Jindra 2019; Truman 2019; Almudi et al. 2020). Since functional wings cannot moult, the winged stage must be the terminal stage, leading to the evolution of an accelerated suite of changes in the final metamorphic moult. Ephemeroptera presents a notable exception, having a short-lived subimago adult stage with thick, hairy wings, followed by a final moult to adults with smooth and thin wings (Truman 2019; Kamsoi et al. 2021). Genetic control of all these transitions to adulthood, including the special case of the mayflies, relies largely on the differential regulation of a common network of transcription factors and hormonal signals (Jindra 2019; Truman 2019; Thomas et al. 2020; Kamsoi et al. 2021; Okude et al. 2022; Chafino et al. 2023; Campli et al. 2024).

In Crustacea, life histories exhibit a diversity that has been described by a rich body of empirical and microscopy evidence, focusing on characterisation of morphological features (Snodgrass 1956; Martin et al. 2014; Olesen 2018; Haug 2020b). Some crustacean taxa hatch as juveniles fully resembling the adult form, as in the orders Diplostraca, Amphipoda, and Isopoda. Several other lineages hatch as a nauplius, an ovoid larva equipped with three pairs of appendages and one eye (Martin et al. 2014). The nauplius might develop further either by progressive segment addition (anamorphosis) and gradual changes as in Anostraca, or by metamorphosis, such as in the clades Thecostraca, Copepoda, and Euphausiacea (Martin et al. 2014). In these lineages, the nauplius transforms into a morphologically distinct larva, known as the cyprid, copepodid, and furcilia larva, respectively, which represent the stage mediating the transition towards acquisition of adult-specific structures (Martin et al. 2014). For example, the cyprid initiates substrate settlement in barnacles, while the copepodid and the furcilia develop fully functional pairs of swimming legs specific to copepod and krill species (Martin et al. 2014; Bernot et al. 2022). The large order Decapoda is also characterised by metamorphic development, where crabs and shrimps hatch as zoeae, swimming using natatory setae, which moult into the decapodid (megalopa), serving as the transitional phase from the planktonic to benthic lifestyles (Martin et al. 2014; Spitzner et al. 2018). Although generally less-well characterised than for Insecta, genetic control of development to adulthood is, as in insects, largely orchestrated through a common network of transcription factors (Ventura et al. 2013; Sun and Patel 2019; Campli et al. 2024; Bentley et al. 2026).

Pancrustaceans therefore present an exemplary model system to study the repeated major life history transition to metamorphic development. Investigating the molecular basis of these evolutionary transitions is increasingly facilitated by genomic sampling of diverse lineages where metamorphosis has been systematically characterised by empirical and experimental methods. This provides the opportunity to apply comparative phylogenomic methods to investigate the mechanisms underlying pancrustacean life history evolution through metamorphosis. Such approaches have proved powerful in unveiling the genetic foundations of ecological transitions from water to land and from terrestrial surfaces to subterranean caves (Balart-García et al. 2023; Wei et al. 2026). Here we explore gene family evolutionary dynamics potentially associated with life history transitions to metamorphic development in Pancrustacea, by using a dataset of 54 species spanning 26 orders and four metamorphic lineages serving as independent evolutionary replicates. Using the shared framework to define metamorphic lineages, we explicitly disregard the diversity of larval types and early developmental modes and focus only on the presence or absence of a metamorphic progression towards the adult stage. We contrast the inferred gene repertoires associated with each evolutionary transition to metamorphic development with those of reconstructed ancestral nodes and non-metamorphic sister lineages. We identify gene family evolutionary dynamics associated with repeated transitions to metamorphic development. Although the expanding gene families are largely distinct amongst lineages, they repeatedly converge on biological functions associated with development and the physiology of metamorphosis.

## Results

### Phylogenomic sampling of pancrustacean diversity enables comparisons of independent transitions to metamorphic development

To compare gene family evolution in metamorphic and non-metamorphic lineages diverging since the Pancrustacea last common ancestor (LCA), a dataset comprising 54 representative species was built by selecting high-quality publicly available genome annotations (see Materials and Methods). From the insects, a maximum of two species per order were selected to achieve a balanced taxonomic distribution, prioritising those with more extensive bodies of supporting literature. Beyond Insecta, species with high-quality annotated gene sets were selected to maximise the representation of pancrustacean developmental diversity. A time-calibrated species phylogeny was estimated for the subphylum-wide dataset that represents 26 pancrustacean orders (Figure 1, Supplementary Figure 1, Supplementary Tables 1, 2, and 3), and which is in agreement with previous large-scale phylogenomics studies (Bernot et al. 2023). Within the phylogeny, four clades are characterised by metamorphic development based on the framework defined above (Metamorphosis LCAs, Supplementary Table 4): Insecta, including orders from Ephemeroptera (mayflies) to Diptera (flies); Eucarida, with the orders Decapoda (prawns, shrimp, lobsters, and crabs) and Euphausiacea (krill); Copepoda, planktonic and benthic copepods with the orders Calanoida and Harpacticoida; and Thecostraca, with the barnacle orders Balanomorpha and Pollicipedomorpha. The LCAs of these clades can be contrasted with two sets of nodes that do not exhibit metamorphic development: the LCAs of each of the four sister non-metamorphic clades of the metamorphic nodes (Sister LCAs, Supplementary Table 4), and the six ancestral nodes up to the Pancrustacea root (Deep nodes). The Sister LCA nodes comprise the non-Insecta hexapods, with the orders Symphypleona (globular springtails) and Entomobryomorpha (springtails), as sister to Insecta; Peracarida, with the orders Isopoda and Amphipoda, as sister to Eucarida; Branchiopoda, with the orders Anostraca (fairy shrimp) and Diplostraca (water fleas), as sister to Copepoda; and Podocopida (ostracods), to contrast Malacostraca and Thecostraca. The Deep nodes represent ancestors that most likely displayed non-metamorphic, anamorphic development of post-embryonic segment addition (Martin et al. 2014; Chipman 2025). The sets of Metamorphosis LCAs and their Sister LCAs are not statistically different in terms of when in evolutionary time they appeared, or in terms of the millions of years separating them from their closest ancestor nodes (Wilcoxon-Mann-Whitney Test with permutation, Age of appearance: p=0.39, Closest ancestor: p=0.56) (Supplementary Figure 2). The dataset therefore enables comparisons across multiple metamorphic and non-metamorphic pancrustacean nodes, which are hereafter referred to as Metamorphosis LCAs, Sister LCAs, and Deep nodes, with Younger nodes referring to the descendant nodes of Metamorphosis LCAs and Sister LCAs.

**Figure 1.**
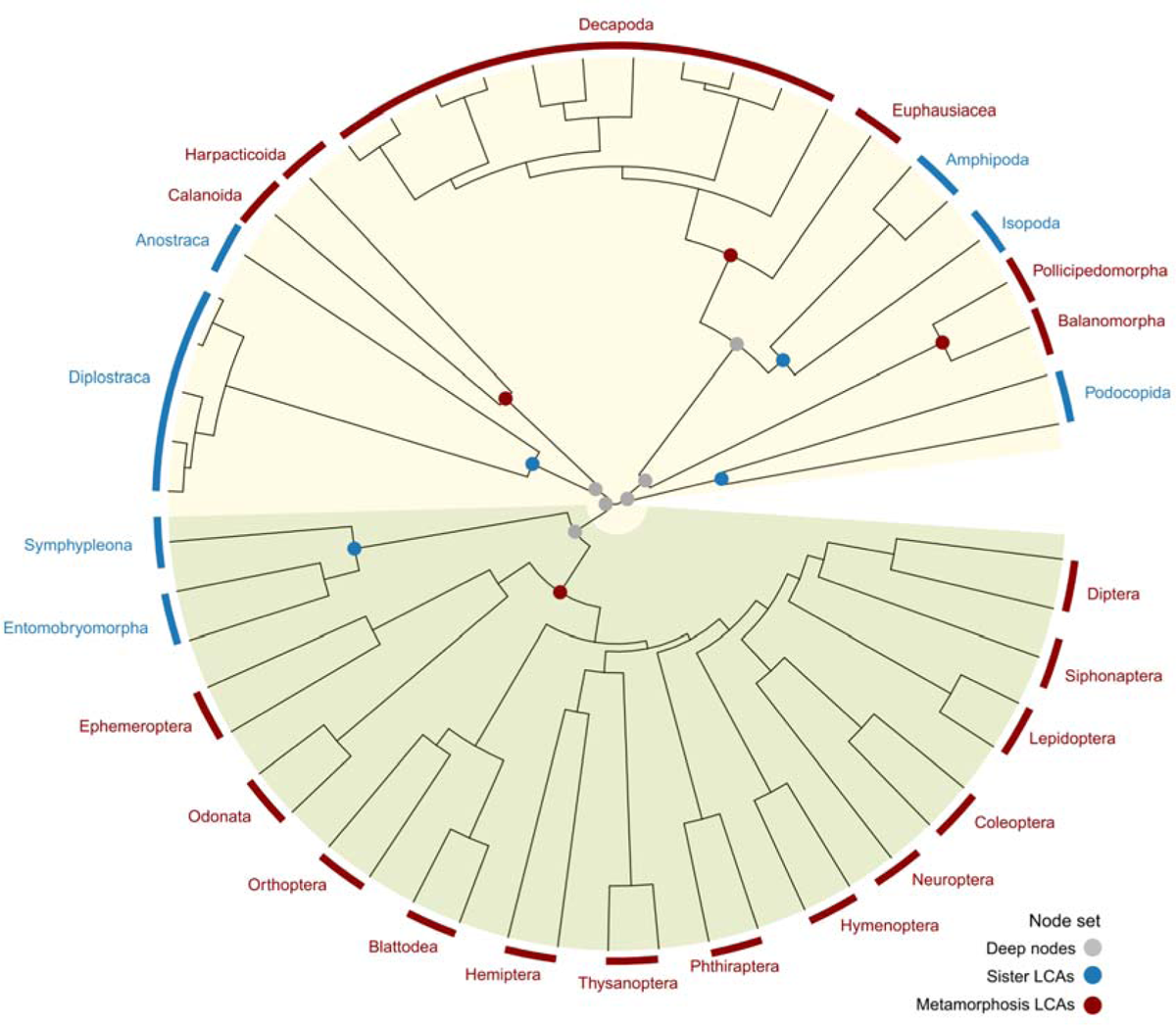
Pancrustacean phylogenomic investigations of gene family evolution associated with transitions to metamorphic development. The pancrustacean phylogeny of the complete dataset of 54 arthropod species, labelled with the 26 represented orders. The orders and key internal nodes are annotated according to their developmental mode: metamorphosis last common ancestors (LCAs) (red), non-metamorphic sister LCA nodes (blue), and deep nodes (grey). Green background: Hexapoda; yellow background: other Pancrustacea. The phylogeny was estimated using a super-alignment of protein sequences from 121 single-copy orthologues and is consistent with previous large-scale phylogenomics studies (Bernot et al. 2023).

### Independent transitions to metamorphic development are characterised by elevated numbers of gene family births and expansions

To quantify gene family evolution across the phylogeny, orthologous groups (OGs) at the Pancrustacea LCA were delineated (see Materials and Methods). The orthology dataset of 42’841 OGs includes 73% of all protein-coding genes from the 54 analysed species (Supplementary Figure 1). Phyletic age assignment classified OGs according to the LCA of all species with orthologues in the OG, distinguishing gene families likely present in the pancrustacean ancestor from those that arose during the diversification of the represented lineages. A majority of 42.2% of all OGs are younger than the Metamorphosis and Sister LCAs, having emerged at one of the 38 descendant Younger nodes (Supplementary Figure 3). A large fraction of OGs date to the pancrustacean ancestor (37.3%), with a further 8.8% emerging at one of the six Deep nodes leading to the Metamorphosis and Sister LCA nodes. OGs that emerged at one of the four Metamorphosis LCAs comprise 8.7% of the total (3’722 OGs), distributed amongst all four LCAs (Figure 2A, “Births”), which is in stark contrast to the much lower fraction found to have emerged at the Sister LCAs (2.6%, 1’112 OGs). Given that there is no statistical difference between Metamorphosis LCAs and Sister LCAs in terms of their ages of appearance, or in terms of the times elapsed since their closest ancestor nodes, this implies a remarkably elevated number of gene family births at the Metamorphosis LCAs. As well as these emergent OGs being more numerous at the Metamorphosis LCAs, they also exhibit significantly lower levels of sequence divergence than those emerging at the Sister LCAs (Figure 2B, “Births”). These sequence-level constraints are quantified as the evolutionary rate of protein sequence divergence amongst OG member genes, distinguishing families with highly conserved or highly divergent sequences (Supplementary Figure 4). Taken together, the Metamorphosis LCAs harbour substantially more gene family births than the Sister LCAs, and these emergent gene families appear more constrained with lower levels of sequence divergence. Transitions to metamorphic development are therefore associated with an increase in births of relatively conserved gene families.

**Figure 2.**
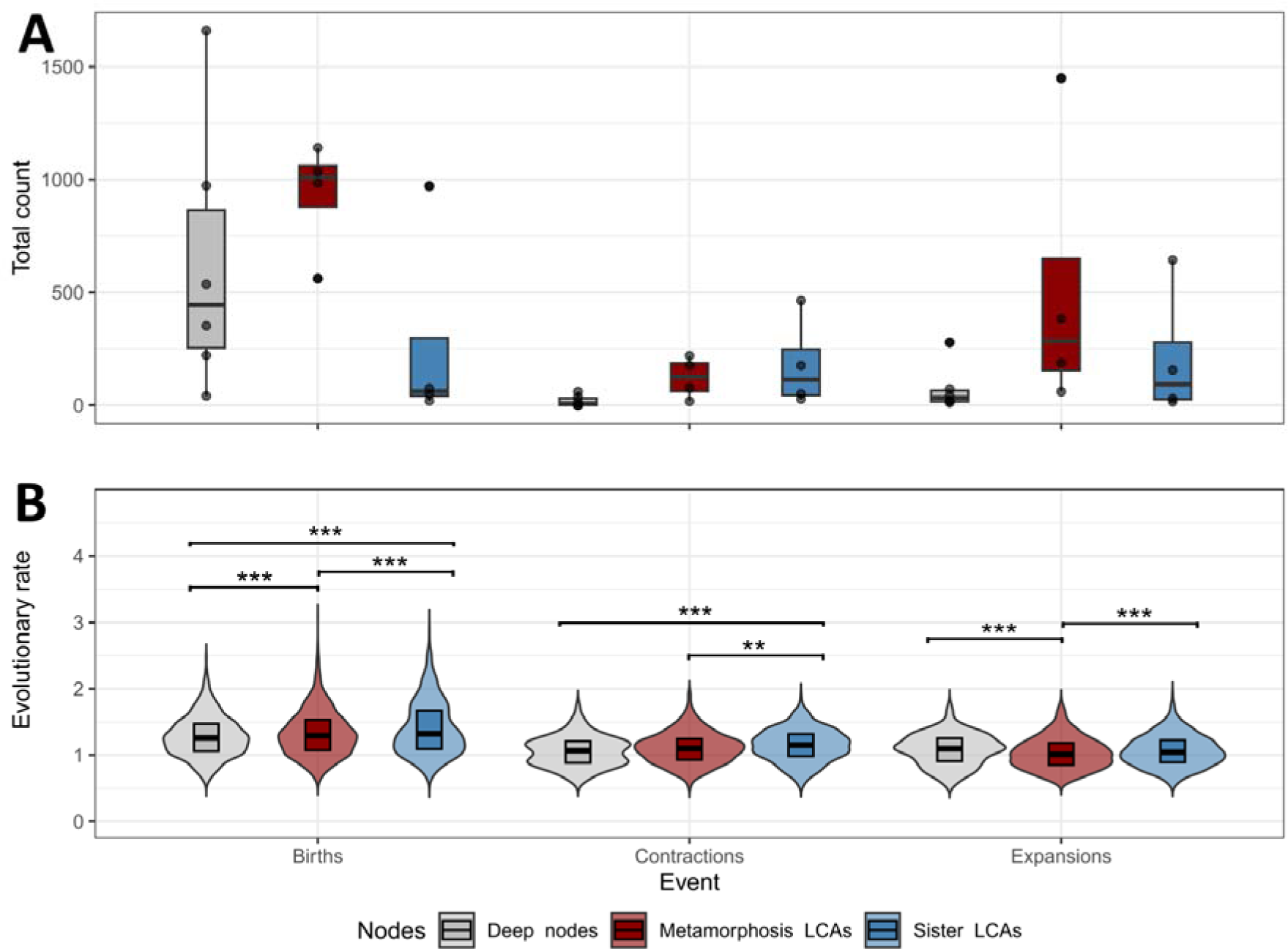
Gene family evolutionary dynamics at the Metamorphosis LCAs, Sister LCAs, and Deep nodes across Pancrustacea. (A) Total counts of gene family evolutionary events inferred to have occurred at the Metamorphosis LCAs, Sister LCAs, and Deep nodes of interest. Each data point shows the total count of orthologous groups (OGs) emerging (births, gene families inferred to originate at that node), decreasing in gene copy number (contractions, phyletic age of the family is the Pancrustacea ancestor), or increasing in gene copy number (expansions, phyletic age of the family is the Pancrustacea ancestor), at each node set. The boxplots show the median, first and third quartiles, and lower and upper extremes of the distribution (1.5 × Interquartile range). (B) Distribution comparisons of OG average inter-species pairwise protein sequence divergence, normalised to the average of single-copy orthologues. Evolutionary rates (y axis) are compared amongst the sets of OGs that are classified as emerging (births), decreasing (contractions), or increasing (expansions) at each set of nodes of interest. Asterisks indicate Wilcoxon test p-values of *** p<0.0005; ** p<0.005, and *p<0.05. The violin plots show the smoothed kernel densities of the distributions, and the boxplots within show the median as well as the first and third quartiles of the distributions.

Filtering the orthology dataset to retain OGs present in the Pancrustacea LCA and with orthologues in at least 85% of the species, ancestral state reconstruction was performed to infer occurrences of gene gains (OG expansions) and losses (OG contractions) across the phylogeny. Amongst these ancient and widespread OGs, a total of 6’919 expansion and 4’186 contraction events were identified across the 53 internal nodes of the pancrustacean tree (Supplementary Figure 3). Of these, 2’078 OGs showed gene gains at the Metamorphosis LCAs, while only 843 and 442 OGs experienced gains mapped to the Sister LCAs and Deep nodes, respectively (Figure 2A). Again, as the Metamorphosis LCAs and Sister LCAs are comparable in terms of their ages and their closest ancestor nodes, these mapped gene gain events indicate an elevated number of gene family expansions at the Metamorphosis LCAs, although the magnitude of the difference is driven largely by the high number found for the barnacles. With respect to gene losses, the Sister LCAs showed the overall highest number with a total of 717 OGs affected, in contrast to just 488 and 110 OGs that experienced losses at the Metamorphosis LCAs and Deep nodes, respectively (Figure 2A). Contrasting sequence-level constraints of these OGs shows that for both expansions and contractions the OGs at the Metamorphosis LCAs exhibit significantly lower levels of sequence divergence than those at the Sister LCAs (Figure 2B). Therefore, as observed for gene family births, there are overall more expanding gene families at the Metamorphosis LCAs than the Sister LCAs, and they also appear to be more constrained. Transitions to metamorphic development are thus also characterised by a proliferation of expansions of ancient and relatively conserved gene families.

### Independent transitions to metamorphic development show convergent functional signatures of gene family evolution

To investigate the functional properties of gene families emerging, expanding, or contracting at the key nodes of interest, OGs were annotated with the Gene Ontology (GO) terms of their member genes from GenBank, RefSeq, FlyBase, and InterPro (see Materials and Methods). Although the emerging OGs were too sparsely annotated to identify GO terms shared across all four Metamorphosis LCAs, contrasting all gene families that emerged at any of the Metamorphosis LCAs with births at any Sister LCAs or Younger nodes identified enrichments for 70 biological processes including “metamorphosis”, “cellular component assembly involved in morphogenesis”, “developmental growth involved in morphogenesis”, “regulation of anatomical structure morphogenesis”, “compound eye morphogenesis”, “chitin-based cuticle development”, “post-embryonic animal organ development”, “activation of immune response”, and “axon guidance” (Supplementary Figure 5, Supplementary Table 5). The same contrast for gene family births at Sister LCAs found only 23 enriched GO terms, with several related to metabolic processes but none with clear links to development or morphogenesis (Supplementary Table 6).

Being widespread and ancient, the gene families expanding and contracting at the nodes of interest were much more richly annotated than the emerging OGs, enabling robust comparisons of results from GO term enrichment analyses. Contrasting gene families that changed in size with those showing stable copy numbers at each node identified sets of enriched GO terms that were then compared to identify common functions characterising the four Metamorphosis LCAs, the four Sister LCAs, or the six Deep nodes. Of the 100 unique GO terms enriched amongst gene families expanding at any of the Metamorphosis LCAs, 60 are common to all four nodes. Semantic similarity clustering of these shared enriched terms summarises the functional characterisation of these gene families in a reduced set of 28 clusters of related GO terms (Figure 3A, Supplementary Table 7). These include a set of biological processes related to the nervous system and development: “central nervous system development”, “morphogenesis of an epithelium”, “developmental maturation”, and “chitin-based cuticle development”. Another set is related to signalling and regulation such as “positive regulation of cell population proliferation”, “regulation of cell differentiation”, “neuropeptide signalling pathway”, and “regulation of autophagy”. A third, smaller set is implicated in membrane (re)organisation: “plasma membrane invagination”, “actin cytoskeleton organisation”, and “synapse assembly”. Others relate to interactions with the external environment “adult feeding behaviour” and “defense response”, or additional development-related processes such as “trunk segmentation” or “imaginal disc growth”. Notably, the remarkably large proportion of GO terms found in common is not due to independent, parallel expansions within the same OGs at all four Metamorphosis LCAs, as the majority of expansions are not shared amongst nodes (Supplementary Figure 6). The identified gene family expansion events involve mostly distinct genes, which nevertheless are annotated with similar biological functions, primarily related to development and other processes relevant in the context of the physiology of metamorphosis. Thus, the repeated transitions to metamorphic development are characterised not by parallel expansions of the same gene families, but by expansions of distinct gene families that converge on similar biological functions.

**Figure 3.**
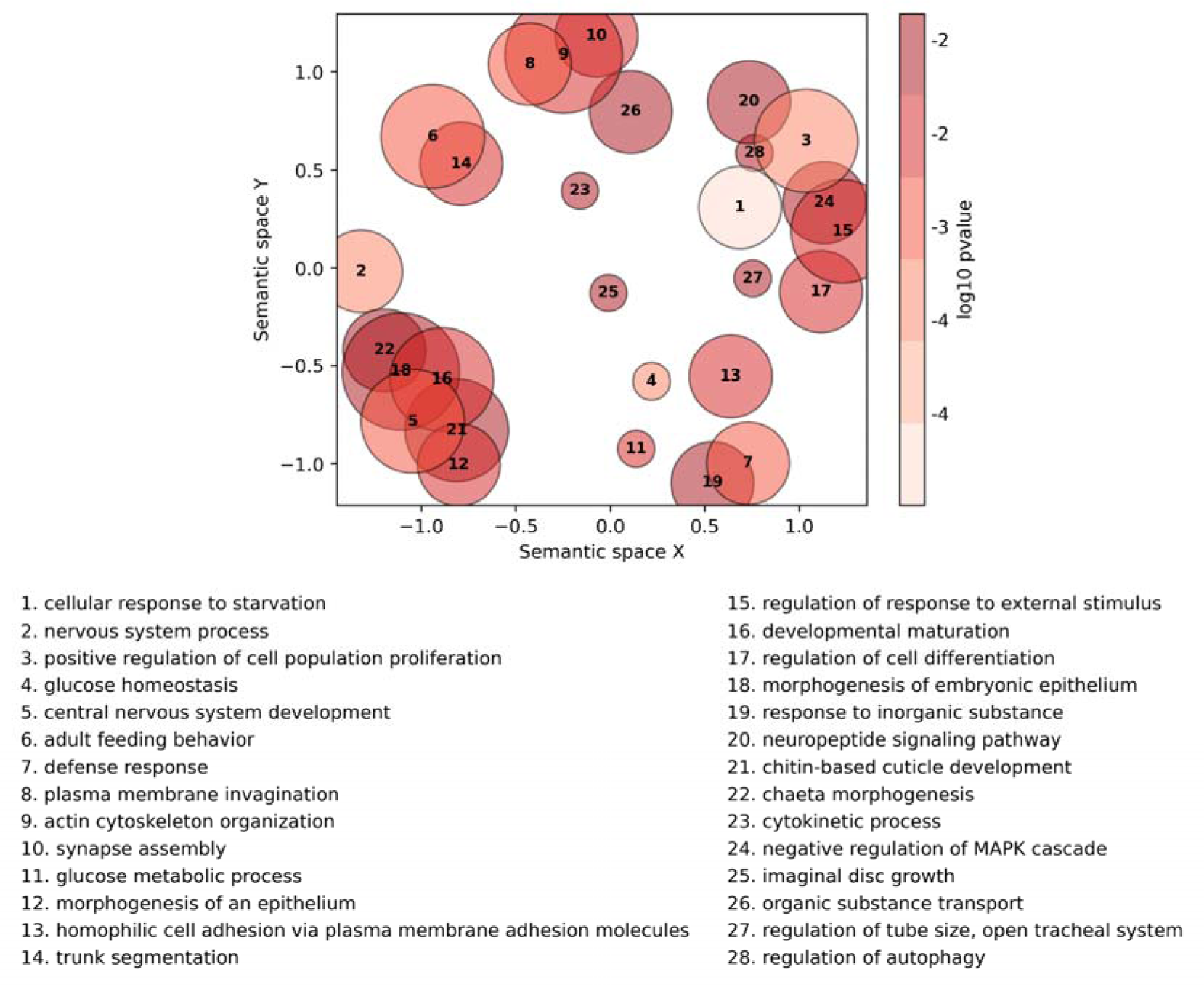
Semantic similarity plot summarising biological functions enriched amongst expanding gene families and shared across Metamorphosis LCAs. Semantic similarity clustering of the 60 Gene Ontology (GO) terms found in all four Metamorphosis LCAs to be enriched amongst gene families with expansions. The circles represent clusters of semantically similar GO terms, based on their relative frequencies in the GO Annotation of UniProt. Larger circles group a higher number of closely related terms together in the same semantic space, the smallest circles representing a single term each. The positioning of the circles along the X and Y axes maximises functional distinctions in the semantic space, where more similar functions are closer together on the plot. Circle colours represent significance values (log10 p-value) from the GO enrichment analysis, according to the legend. Number labels on each circle correspond to the presented list of GO term names, ordered by significance values.

In stark contrast to the Metamorphosis LCAs, sets of GO terms enriched amongst the gene families expanding at the other nodes of interest have little in common, with very few shared by all Sister LCAs (n=12) or all Deep nodes (n=6) (Supplementary Tables 8 and 9), and with terms that are generally less specific than those found for the Metamorphosis LCAs, such as “multicellular organismal process”, “transmembrane transport” or “proteolysis”. The smaller sets of families showing contractions at the nodes of interest are characterised by few enriched GO terms, with 81, 35, and 49 unique terms for Metamorphosis LCAs, Sister LCAs, and Deep nodes, respectively. Term list comparisons show there are very few in common for the Metamorphosis LCAs (n=18, Supplementary Table 10), or for the Sister LCAs (n=6, Supplementary Table 11), and none for the Deep nodes. As for the expanding gene families with common terms across the Sister LCAs or the Deep nodes, the few identified GO terms refer to rather general processes such as “carbohydrate metabolic process” or “proteolysis”. Considering these node comparisons for expansion or contraction events, the high degree of overlapping enriched GO terms, and the associations of these terms with specific processes conceivably relevant for the biology of metamorphosis, are thus features uniquely characterising the gene families expanding at the Metamorphosis LCAs.

### Gene families exhibiting adaptive, lineage-specific expansions include genes implicated in neural and sensory development, morphogenesis, and moulting

To determine if some expansions were associated with adaptive evolution, all 528 expanding gene families with exact-matching enriched GO terms shared across the Metamorphosis LCAs were tested to assess whether increases in family sizes showed evidence of lineage-specific adaptive signal (Beaulieu et al. 2012; Seppey et al. 2019). The Brownian motion (BM) and the Ornstein-Uhlenbeck (OU) models were used to describe evolutionary scenarios under neutral and selective pressures. In the first case, gene copy numbers change across lineages characterised by metamorphic and non-metamorphic development solely due to stochastic processes (null hypothesis). In the second case, gene family sizes evolve driven by selective pressures to reach an “optimum” value, which may (null hypothesis) or may not (alternative hypothesis) differ amongst lineages (see Materials and Methods). In total, 15 families (3%) tested positive for selection towards two optima (delta mean copy number > 0.7, delta AICc > 2, delta optima > 1), where the optimum for the Metamorphosis LCAs was the largest (Figure 4A). The 22 shared enriched GO terms annotated to the 15 gene families include processes related to development (“nervous system development” and “chitin-based cuticle development”), morphogenesis (“dorsal closure”, “head involution”, as well as “salivary gland”, “chaeta”, and “imaginal disc-derived wing vein morphogenesis”), and regulation of cell proliferation, of gene expression, and of Mitogen-Activated Protein Kinase (MAPK) signalling (Supplementary Tables 12 and 13).

**Figure 4.**
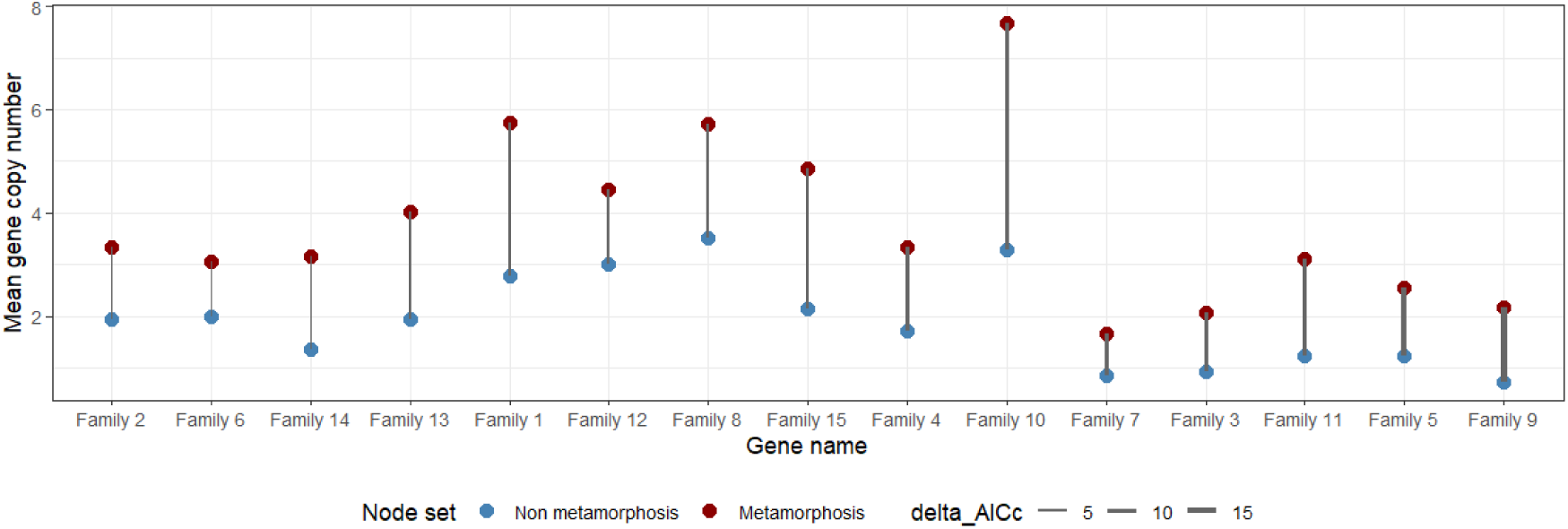
Signatures of adaptive lineage-specific gene family expansions at the Metamorphosis LCAs. Gene families annotated with enriched Gene Ontology (GO) terms found across all the Metamorphosis LCAs for which modelling evolution by selection (H1: OU models) for higher gene copy numbers (y-axis) in metamorphic species than non-metamorphic species showed a greater likelihood than stochastic neutral evolution (H0: Brownian models). Model support is indicated by differences in the Akaike Information Criterion corrected for small sample size (AICc) values (line width). Inclusion cut-offs were set as having a difference in AICc greater than 2 and a difference in mean copy number greater than 0.7. Gene family GO term annotations and protein identifiers of genes from *D. melanogaster* are presented in Supplementary Table 13.

Amongst the gene families that tested positive for signatures of adaptive lineage-specific gene family expansions at the Metamorphosis LCAs are the transcription factors encoded by *knirps* (Family 11) and *hairy* (Family 5). These are transcriptional repressor proteins which are involved in both embryonic and neural development, regulating segmentation by controlling pair-rule genes (San-Juán and Baonza 2011; Yamasaki et al. 2011; Urbach et al. 2016; An et al. 2017; Clark et al. 2019; Sato et al. 2019; Myasnikova and Spirov 2020). Other expanding gene families have also been implicated in neurogenesis processes: the immunoglobulin adhesion proteins of the *klingon* subfamily (Family 1) (Cheng et al. 2019; Shimozono et al. 2019), the opsin-binding *arrestin* gene family including the nonvisual Kurtz arrestin (Family 12) that acts an inhibitor of MAPK and Toll pathways during development (Shieh et al. 2014; Nolan et al. 2024), the *inscuteable* family (Family 7) of adaptor proteins that regulate cell division particularly for neuroblasts and epithelial cells (Culurgioni et al. 2011; An et al. 2017), the transmembrane proteins from the *tartan*-like family (Family 3) implicated in neuronal cell and tracheal network development (Krause et al. 2006; Ylla et al. 2018; Wang and Dahmann 2020; Wang and Dahmann 2020), and the *kekkon*-like family (Family 9) with roles in neuronal differentiation (MacLaren et al. 2004; Ulian-Benitez et al. 2017). The immunoglobulin subfamily including the *Drosophila* gene *klingon* mediates cell adhesion and is involved in development of photoreceptor neurons (Cheng et al. 2019; Shimozono et al. 2019). The transmembrane leucine-rich repeat protein Tartan is also involved in axon guidance, along with tracheal morphogenesis and establishment of the dorso-ventral boundary in imaginal wing discs, by binding its receptor transmembrane leucine-rich receptor Capricious (Krause et al. 2006; Wang and Dahmann 2020; Wang and Dahmann 2020). Harbouring both transmembrane leucine-rich and immunoglobulin-like domains, proteins of the *kekkon* family have been shown to contribute to synaptic growth, together with Toll receptors (MacLaren et al. 2004; Ulian-Benitez et al. 2017). They are also inhibitors of the Epidermal Growth Factor Receptor (EGFR) pathway in developing imaginal eye and wing discs (MacLaren et al. 2004; Ulian-Benitez et al. 2017). Arrestins also play ancient and conserved roles within the photoreceptor, where proteins encoded by paralogous genes have distinct rhodopsin regulatory functions (Shieh et al. 2014; Nolan et al. 2024). Almost all the knowledge about the functions of these genes comes from studies in model insects, and their functional evolution in Pancrustacea may have been shaped by pleiotropy, regulatory rewiring, and co-option or exaptation. Nonetheless, both the known function and the evolutionary patterns of these pathways and processes appear relevant to the biology and evolution of metamorphosis.

## Discussion

The growing availability of annotated genomes for non-insect arthropods enabled the construction of a phylogenomic dataset to investigate the underlying molecular foundations across four independent transitions to metamorphic development. The large arthropod clade Pancrustacea includes a great diversity of developmental modes, often characterised by lineage-restricted life-stage morphologies (Snodgrass 1956; Bishop et al. 2006; Martin et al. 2014; Spitzner et al. 2018; Jindra 2019; Truman 2019; Haug 2020b; Vea and Minakuchi 2021). The distinctiveness of such morphologies makes comparisons of life stages complex. Nevertheless, a framework to define metamorphic life-stage progressions facilitates integrating the characterisation of pancrustacean development with the exploration of broad genomic patterns, employing an operational flexibility emerging in the research community (Bishop et al. 2006; Martin et al. 2014; Spitzner et al. 2018; Jindra 2019; Truman 2019; Vea and Minakuchi 2021). We define metamorphosis as being characterised by major morphological and ecological changes coupled with the progression towards the adult stages. Our framework builds on accumulated morphological knowledge to define metamorphic developmental progressions independently of the homology of individual life stages. For example, a larval-pupal metamorphosis is not assumed to be homologous to a zoea-megalopa metamorphosis, despite both being recognised as metamorphic life-stage progressions based on lineage-specific morphological characters (Martin et al. 2014; Spitzner et al. 2018; Truman 2019). Together, the phylogenomic dataset and the generalised framework of metamorphosis facilitate the integrative exploration of gene family evolution in the context of repeated major life history transitions to metamorphic development across Pancrustacea.

Exploring gene family evolutionary dynamics associated with repeated transitions to metamorphic development in Pancrustacea reveals several key patterns. (1) The Metamorphosis LCAs exhibit more gene family births and more gene family expansions compared to the Sister LCAs. Both these sets of emerging and expanding gene families appear to be more constrained, with lower levels of sequence divergence than those of the Sister LCAs. (2) Gene families emerging at the Metamorphosis LCAs are enriched for functions related to development and morphogenesis, while those of the Sister LCAs are not. (3) Gene families expanding at the Metamorphosis LCAs show a remarkably high overlap amongst their enriched functional annotations, which relate to developmental and other processes relevant to the physiology of metamorphosis; those of the Sister LCAs and Deep nodes, as well as for gene family contractions for any set of nodes of interest, do not. (4) In spite of their common functions, the expansions at the Metamorphosis LCAs involve mostly distinct genes rather than parallel expansions within the same gene family at all four nodes. This striking pattern points to functional convergence through distinct genetic trajectories underpinning repeated transitions to metamorphic development across Pancrustacea. (5) Despite mostly involving distinct genes, signatures of adaptive lineage-specific gene family expansions at the Metamorphosis LCAs are evident for a small subset, which includes genes implicated in pathways and processes associated with metamorphic development.

These observed patterns are based on the analyses of a phylogenomic dataset that is broad and representative of Pancrustacea, with 54 species from 26 orders. However, the dataset is limited in the depth of representation across the non-insect lineages. For example, availability of high-quality annotations from additional non-metamorphic lineages is currently limited, such as the ametabolous bristletails and silver fishes from the insect orders Archaeognatha and Zygentoma, and the centipede-like crustaceans from the class Remipedia. As taxonomic representation can impact orthology delineation and ancestral state reconstructions, some of the inferred events characterising gene family dynamics across the phylogeny may be less well supported than others. Greater and more diverse species sampling within the lineages of interest would enhance the confidence of inferred gene family dynamics at key internal nodes, allowing more robust comparisons of the Metamorphosis LCAs with the Sister LCAs and Deep nodes. For example, Cirripedia are represented by only two single-species orders in the dataset, but the barnacles include almost ten recognised taxonomic orders, and all of them develop through the cypris larva, harbouring four pairs of antennules specialised to the attachment phase on benthic substrates or on the host (Martin et al. 2014; Bernot et al. 2022). The barnacles present an additional complexity as they might have experienced an ancient whole genome duplication (WGD) event (Kim et al. 2019; Bernot et al. 2022; Yuan et al. 2024), with differential gene loss or maintenance post WGD. This lineage-specific feature could contribute to the larger number of expansions identified at the Thecostraca LCA. The caveats arising from biased taxonomic sampling could impact inferences at all nodes of interest. Nevertheless, the enrichments of functions related to development or morphogenesis, and the large overlap of enriched processes, are unique to the Metamorphosis LCAs. This suggests that the observed patterns are underpinned by a signal that is robust, despite the limited depth of representation beyond insects. Fortunately, improved representation across all groups in the dataset to match the depth of coverage within the insects may be achievable in the future with the generation of additional high-quality genomes and annotations (Mazzoni et al. 2023; Blaxter et al. 2025).

At the level of individual gene families whose evolutionary dynamics are linked to life history transitions to metamorphic development, most functional information is derived from model insects. This is evident from the observed paucity of functional annotations for lineage-specific emergent gene families, which could represent important novelties with functional roles that remain to be elucidated. Where extrapolations from better-studied models to other lineages are possible, they must remain cautious and context-aware, especially given the dynamic changes in gene copy number of these families. Knowledge of the pathways and processes in which they play a role provides useful guidance, but it is important to consider pleiotropic effects, as well as post-duplication regulatory rewiring and pathway co-option (True and Carroll 2002). Nonetheless, the involvement of several genes from families involved in embryonic and nervous system development, segmentation, neurogenesis, developmental MAPK, EGFR, and Toll pathway signalling, tracheal and epithelial morphogenesis, and photoreceptor development, points to plausible sources of evolutionary changes that supported transitions to metamorphic development. Pair-rule genes like *hairy* in general have a conserved role in the early stages of the segmentation cascade across arthropods, although the details vary amongst different lineages (Janssen et al. 2012; Green and Akam 2013; Auman and Chipman 2018; Clark et al. 2019; Reding et al. 2019; Chipman 2025). Whereas gap genes like *knirps* have a much less conserved role in segmentation, probably only being relevant in insects (Chipman 2020), where *knirps* itself may play a Diptera-specific role in this context (Naggan Perl et al. 2013). Interestingly, *knirps* and *hairy* also regulate expression of several moult-inducing ecdysteroid biosynthetic enzymes in prothoracic glands (Okamoto and Yamanaka 2020; Kamiyama and Niwa 2022; Fernandez-Nicolas et al. 2023). Along with *deadpan*, *knirps* and *hairy* are also involved in insect neural development (San-Juán and Baonza 2011; Yamasaki et al. 2011; Urbach et al. 2016; An et al. 2017; Sato et al. 2019; Myasnikova and Spirov 2020). Participating in both processes is not uncommon, as several other genes are known to be firstly expressed during embryonic patterning and subsequently acting again during neurogenesis and moulting (Truman 2019; Chafino et al. 2023; Truman et al. 2023; Wexler et al. 2023; Alon and Chipman 2025). Neural morphogenesis is also controlled by *tartan* and *capricious*, which mediate cell interactions, defining boundaries between cell populations, particularly in the wings, eyes, and nervous system (Milán et al. 2001; Krause et al. 2006). Similarly in the development of photoreceptor neurons, *klingon* mediates cell adhesion processes, as well as axonal guidance, and other neuronal wiring-related functions (Cheng et al. 2019; Shimozono et al. 2019). Such cell sorting activities and morphogenic regulatory functions would likely be particularly important in metamorphic morphological reorganisations to reach the adult form. Transitions in Pancrustacea might therefore be based on the recruitment of genes from pleiotropic families that can act at different levels during development, such as during segmental patterning, neurogenesis, and moulting.

The links to functions associated with neural development and photoreception may also reflect the adaptive landscape changes through metamorphosis. Complex neural structures are required to integrate abiotic and biotic stimuli from the environment, and responses to light stimuli play an important role in processes specific to adulthood, such as partner recognition mechanisms or changes in feeding strategies. For example, mayflies harbour an expanded toolkit for photoreception, with male-specific eye structures, and aquatic juveniles and aerial adults display different gene expression profiles, showing a shift from odorant-based to vision-based sensation (Almudi et al. 2020). Compound eyes of some dragonflies grow entire new regions during the last moult (Chou et al. 2020). Mantis shrimps have compound eyes highly specialised for predation, with optic neuropils that develop during the last moult at the onset of adulthood and totally replace degenerating larval optic lobes, similarly to the visual system remodelling observed in holometabolous insects (Chou et al. 2020). Other functions with plausible links to adaptive landscape changes include immune responses and cuticle/epithelial remodelling. In both insects and crustaceans, the immune system is activated during moulting, likely as a stress response and protection against pathogens (Gao et al. 2015; Nunes et al. 2021; Liu et al. 2022; Betancourt et al. 2024; Kim et al. 2026). A protective role is also played by the exoskeleton at the interface with the environment, which is composed of layers of chitin fibres and cuticle proteins secreted by underlying epithelial cells, and whose overall structure is conserved in Pancrustacea (Roer et al. 2015). Specific exoskeleton structures, however, vary considerably, with chaetae or setae and their associated neural projections showing morphologies, and likely their underlying gene expression programmes, that are highly lineage-specific (Mellon 2012; Bello et al. 2022; Kelly et al. 2023; Kor et al. 2023; Kim et al. 2026). While maintaining a cautious approach to functional extrapolations, the results point to a plausible interpretation where multi-phasic complex lifestyles might convergently involve common functional processes, despite differences in the specific adaptations that mediate their life history transitions, supporting a hypothesis of convergence between Insecta and Crustacea. These observations are consistent with the adaptive landscape changes that define metamorphic development under our framework.

While plausibly relevant to the physiology of metamorphosis, the functions of gene families with evolutionary dynamics associated with transitions to metamorphic development are not necessarily unique to this particular life history transition. Recent investigations of gene family turnover dynamics and the transition to life on land have also revealed links with common processes including metabolism and reproduction, immunity, embryogenesis, development, and cell differentiation and morphogenesis (Benítez-Álvarez et al. 2026), as well as with exoskeleton structure, water conservation, and sensory development (Wei et al. 2026). Expansions were also more prominent than contractions, and specific gene families related to exoskeleton formation and moulting were also identified amongst the terrestrialisation genomic toolkit (Benítez-Álvarez et al. 2026). Thus, despite transitions to metamorphosis and life on land having clear underlying differences, some functional groups of gene families involved in these processes represent a common toolkit that participates in the reshaping of organismal biology to adapt to abiotic and biotic changes. Importantly, while functional convergence is evident, repeated evolutionary innovation does not require the repeated recruitment of the same genes. These findings therefore collectively reframe the role of moulting in arthropod diversification from a necessary step for proper growth and development to an evolutionarily flexible developmental substrate that can potentiate truly remarkable life history transitions.

## Materials and Methods

### Publicly sourced genome assemblies and annotations

The pancrustacean genome assemblies with annotated proteomes available from the United States National Center for Biotechnology Information (NCBI) were evaluated using the Arthropod Assembly Assessment Catalogue (A3Cat, release 2025-01-01) (Feron and Waterhouse 2022). These were filtered based on the A3Cat Benchmarking Universal Single-Copy Orthologues (BUSCO) scores: the annotated assembly having the highest Arthropoda Complete BUSCO score was retained where more than one assembly was available for individual species; all selected assemblies were required to have scores of less than 30% Arthropoda Duplicated BUSCO and at least 75% Arthropoda Complete BUSCO. These thresholds were applied specifically to avoid the exclusion of some non-insect Pancrustacea from key underrepresented lineages required to ensure a broad representation of pancrustacean diversity. Subsequently, to produce a more balanced phylogenetic representation, insect proteomes were selected by retaining within each taxonomic order a maximum of two species, to include model systems with previous accumulated knowledge and species characterised by neometabolic (Lepidoptera: *Bemisia tabaci*; Thysanoptera: *Frankliniella occidentalis*, *Thrips palmi*) and paedomorphosis (Coleoptera: *Ignelater luminosus*; Blattodea: *Zootermopsis nevadensis*) development (Vea and Minakuchi 2021). Proteomes, annotation feature tables and Gene Ontology (GO) term annotations from the selected assemblies were downloaded from NCBI repositories GenBank and RefSeq (2025-03-17) and assessed for completeness using BUSCO v5.4.3 and the arthropoda_odb10 lineage dataset (option: -mode protein) (Manni et al. 2021). All annotated proteomes with a score of at least 75% Arthropoda Complete BUSCO were retained and filtered for the longest protein-coding isoform based on their corresponding annotation feature table. This resulted in the selection of the annotated proteomes of 54 pancrustacean species representing 26 orders from the two major clades within the paraphyletic Pancrustacea, with Arthropoda BUSCO Complete scores of median 95.8% (mean 93.4%, standard deviation 6.3) and Arthropoda BUSCO Duplicated scores of median 2% (mean 4.6%, standard deviation 7.6), and protein-coding gene counts of median 17’059 (mean 19’806, standard deviation 8’588). Assembly accessions, protein-coding gene counts, and BUSCO assessments are provided in Supplementary Table 1. To augment the NCBI-sourced functional annotations of the proteomes, GO term annotations for *D. melanogaster* were additionally sourced from FlyBase (19/05/2025) (Öztürk-Çolak et al. 2024).

### Species phylogeny reconstruction

The species phylogeny was computed from 121 single-copy orthologues found in at least 95% of the total set of 54 species, detected by the BUSCO assessments performed for the A3Cat on the genome assemblies using the arthropoda_odb10 lineage dataset. Protein sequences were aligned using MUSCLE v3.8.1551 with default parameters and trimmed using the automated method for threshold optimisation in TrimAl v1.4.1 (option -strictplus) (Edgar 2004; Capella-Gutiérrez et al. 2009). A concatenated super-alignment was built from the 121 individual alignments: 51’083 columns, 47’966 distinct patterns, 36’774 parsimony-informative, 6’920 singleton sites, 7’388 constant sites, other metrics computed by Alistat v1.15 (Wong et al. 2020) are available as Supplementary Table 2. This supermatrix was employed for phylogeny reconstruction performed using ModelFinder as implemented in IQ-TREE v2.2.0-beta with 1’000 ultrafast bootstrap samples (option -msub nuclear -B 1000 - bnni (Nguyen et al. 2015; Kalyaanamoorthy et al. 2017; Hoang et al. 2018; Minh et al. 2020). The resulting phylogeny was inspected and manually rooted with NJ-plot. The paraphyletic clade of crustaceans, comprising Hexapoda and Crustacea within the monophyletic Pancrustacea was recovered and placement of main clades is in agreement with previous phylogenomics findings (Bernot et al. 2023). The molecular species tree was time-calibrated using ten calibration nodes (Supplementary Table 3), supplied to the functions makeChronosCalib() and chronos of the ape v5.8 R package (Paradis and Schliep 2019). Divergence time estimates were sourced from the TimeTree v5 database (Kumar et al. 2022). The resulting ultrametric species phylogeny representing 500 million years of pancrustacean evolution with 53 internal nodes was used for ancestral state reconstructions. Branch lengths were compared across node sets using permutation tests as implemented in the coin R v.1.4.3 package (Hothorn et al. 2008). The phylogeny was annotated with species developmental modes: the developmental mode for all Insecta species, given the absence of ametabolous species, was annotated as metamorphic (including all holometaboly, hemimetaboly, and neometaboly), for all non-insect Hexapoda species as non-metamorphic, and for the non-hexapod Crustacea species annotations were based on the Atlas of crustacean larvae (Martin et al. 2014) (Supplementary Table 4).

### Orthology delineation

Orthology was delineated at the level of the Last Common Ancestor (LCA) of all the 54 selected arthropod species, where an Orthologous Group (OG) represents the set of genes descended from a single gene in the Pancrustacea LCA. The OrthoDB standalone pipeline OrthoLoger v3.0.2 was run with default parameters (Kuznetsov et al. 2023), after filtering the proteomes for the single longest protein-coding isoform of each gene. The OrthoLoger algorithm uses all-against-all pairwise protein sequence alignments to identify best reciprocal hits (BRHs) between all genes from each species pair in the dataset. Subsequently, a graph-based clustering procedure starting with BRH triangulation progressively builds OGs aiming to identify and include all genes descended from a single gene in the LCA. The resulting orthology dataset comprised a total of 42’841 OGs and 782’987 genes, representing 73% of the input gene set from the annotated proteomes of the 54 species. OG phyletic ages were computed as the LCA of all the species included in the OG, simplified to stratifications of Metamorphosis LCAs (Insecta, Eucarida, Copepoda, and Thecostraca LCAs), Sister LCAs (non-Insecta hexapods, Peracarida, Branchiopoda, and Ostracoda LCAs), Deep nodes up to but excluding the Pancrustacea LCA (6 nodes), and all remaining Younger nodes (38 nodes). OG evolutionary rates, as defined in OrthoDB (Waterhouse et al. 2013), were computed using OrthoLoger ancillary tools by normalising all interspecific pairwise protein sequence identities within each OG against the mean identity of the corresponding BRHs between each species pair, and then averaging the resulting values across the OG. Values close to 1.0 indicate protein sequence divergence similar to the genome-wide average for single-copy orthologues, while lower and higher values indicate slower (more constrained) and faster (less constrained) sequence evolution, respectively.

### Gene count ancestral state reconstruction

Ancestral state reconstruction aims to estimate the likely state (in this case gene copy numbers) at the internal nodes of the species tree given the observed state at the terminal branches and a model of gene family evolution. To estimate ancestral gene copy numbers, and thus map gene family expansion and contraction events on the species tree, a maximum likelihood approach to the birth-death model of gene family size evolution was employed, as implemented in Computational Analysis of gene Family Evolution (CAFE) v5 (Mendes et al. 2021). By treating the observed gene family counts as distributions rather than definitive observations, CAFE is able to account for annotation errors present in the input gene count data. To target ancient and widespread OGs, only OGs with a phyletic age of the Pancrustacea LCA including at least 85% of the total species in the dataset (46/54) were used for the input gene count table provided to CAFE. To ensure model convergence, four independent runs were performed, always estimating one global lambda for the whole tree, with one k rate category and Poisson root family size distributions. Estimation of lambda is 0.0005, supported by a maximum likelihood score of -lnL 270778.

### Functional enrichment testing

Gene Ontology (GO) enrichment analysis was performed using the TopGO R package (v2.54.0) (Alexa and Rahnenführer 2025) with Fisher’s exact test and ranked by the “weight01” algorithm, limited to the top 100 enriched terms. All OGs were annotated with the unique GO terms of their member proteins. For the emerging OGs (i.e. gene family births, identified as OGs with a phyletic age younger than the root Pancrustacea LCA, Supplementary Figure 3A), the foreground sets for enrichment testing consisted of the OGs with a phyletic age of (i) any of the four Metamorphosis LCAs or (ii) any of the four Sister LCAs. The background sets consisted of all OGs emerging at any Younger nodes (younger than Metamorphosis LCAs and younger than Sister LCAs) plus for (i) OGs emerging at the Sister LCAs and for (ii) OGs emerging at the Metamorphosis LCAs. The contrasts therefore control the treatment of foreground and background sets to comprise only emerging OGs, and only OGs emerging at the nodes of interest and subsequently (Younger nodes), with the exclusion of all Deep node emerging OGs and Pancrustacea LCA OGs. The proportions of emerging OGs annotated with GO terms were generally low, even after adding annotations based on protein domain contents assessed with InterProScan v5.59-91.0 using the curated InterPro2GO mapping (Blum et al. 2025). Therefore, for the four Metamorphosis LCAs and for the four Sister LCAs the combined sets of OGs from each were analysed together. The Biological Process GO terms that were significantly enriched (p<0.05) were summarised and visualised with the GO-Figure! Python package (v1.0.1), to reduce term redundancy by semantic similarity, using a similarity cutoff of 0.3 (Reijnders and Waterhouse 2021). For the expanding or contracting OGs (i.e. gene duplications or losses inferred to have occurred at internal nodes of the phylogeny, Supplementary Figure 3B), only OGs with a phyletic age of the Pancrustacea LCA and that included at least 85% of the total species in the dataset were considered (ancient and widespread OGs). The foreground sets for enrichment testing consisted of the OGs (i) expanding, or (ii) contracting, at each of the nodes of interest (Metamorphosis LCAs, Sister LCAs, and Deep nodes). In each contrast, the background sets consisted of all other ancient and widespread OGs that were found to be stable at the same node of interest (i.e. not changing in copy number). The more comprehensive GO term annotations of these OGs meant that resulting lists of significantly enriched (p<0.05) GO terms could be compared to identify Biological Process GO terms common to all four Metamorphosis LCAs, all four Sister LCAs, or all six Deep nodes. These lists of common GO terms were summarised and visualised with the GO-Figure! Python package (v1.0.1), using a similarity cutoff of 0.3.

### Evolutionary model testing

To distinguish lineage-specific adaptive expansions from background variation in gene family size, OGs with expansions at one or more of the Metamorphosis LCAs, and being annotated with any of the GO terms enriched in all the Metamorphosis LCA contrasts, were explicitly tested for adaptive lineage-specific expansions as implemented in the OUwie R package (v2.10) (Beaulieu et al. 2012), adapting code from (Seppey et al. 2019). This requires all nodes and leaves of the species phylogeny to be annotated with their respective developmental modes, therefore, metamorphic development was extrapolated from all Metamorphosis LCAs to their descendent nodes and species, and all the remaining nodes and species were labelled as non-metamorphic. The developmental modes were considered as two different regimes, to compare the per-species gene copy phylogenetic profiles between the two developmental modes and test whether gene copy number values evolve towards an optimum under selective pressure, which differs between the developmental regimes. Null hypotheses H0 comprises: the Brownian Motion (BM) model with a single rate or multiple rate estimates for each developmental mode, respectively BM1 and BMS, describing neutral evolution, where no selection occurs and differences between modes are due to stochastic processes; the Ornstein-Uhlenbeck model (OU) where selection occurs but both regimes evolve towards the same optimum under OU1, thus no signature of lineage-specific adaptive expansion is found. Alternative hypothesis H1 describes that, under adaptive selection, gene copy numbers reach two different optima having the same variance or two different variances, referred to respectively as the OUM and OUMV models, thus supporting signatures of lineage-specific adaptive expansion in metamorphic clades. The Akaike information criterion corrected for small sample size (AICc) was used to select the best model amongst the BM1, BMS, and OU1 models for H0 and the OUM and OUMV for H1, respectively. OGs having the best H1 model preferred over the best H0 model with AICc larger or equal to 2 and reaching an optimum for the metamorphosis regime larger than the non-metamorphic development regime optimum by at least a factor of 1 were included in the significant results, as having signatures of adaptive gene family expansion specific to the metamorphic lineages.

## Data Availability

The genomic data underlying this article are available from the European Nucleotide Archive (ENA) at https://www.ebi.ac.uk/ena/browser/home, and can be accessed with the accession numbers provided in Supplementary Table 1. The orthology dataset and ancestral state reconstruction data generated as part of this study are available at figshare DOI: https://doi.org/10.6084/m9.figshare.33070619.

## Acknowledgements

The authors thank all members of the Sinergia project “An interdisciplinary study of arthropod moulting: linking genotype, phenotype and life history evolution” for useful discussions that supported the development of this manuscript.

## References

Alexa A, Rahnenführer J. 2025. topGO: Enrichment analysis for gene ontology. https://bioconductor.org/packages/topGO

Almudi I, Vizueta J, Wyatt CDR, de Mendoza A, Marlétaz F, Firbas PN, Feuda R, Masiero G, Medina P, Alcaina-Caro A, et al. 2020. Genomic adaptations to aquatic and aerial life in mayflies and the origin of insect wings. Nat. Commun. 11:2631. 10.1038/s41467-020-16284-8

Alon N, Chipman AD. 2025. Neurogenesis in the trunk and brain of the milkweed bug Oncopeltus fasciatus: insights beyond holometabolan models. Front. Zool. 23:3. 10.1186/s12983-025-00593-z

An H, Ge W, Xi Y, Yang X. 2017. Inscuteable maintains type I neuroblast lineage identity via Numb/Notch signaling in the Drosophila larval brain. J. Genet. Genomics Yi Chuan Xue Bao 44:151–162. 10.1016/j.jgg.2017.02.005

Auman T, Chipman AD. 2018. Growth zone segmentation in the milkweed bug Oncopeltus fasciatus sheds light on the evolution of insect segmentation. BMC Evol. Biol. 18:178. 10.1186/s12862-018-1293-z

Balart-García P, Aristide L, Bradford TM, Beasley-Hall PG, Polak S, Cooper SJB, Fernández R. 2023. Parallel and convergent genomic changes underlie independent subterranean colonization across beetles. Nat. Commun. 14:3842. 10.1038/s41467-023-39603-1

Beaulieu JM, Jhwueng D-C, Boettiger C, O’Meara BC. 2012. Modeling stabilizing selection: Expanding the Ornstein–Uhlenbeck model of adaptive evolution. Evolution 66:2369– 2383. 10.1111/j.1558-5646.2012.01619.x

Bello E, Chen Y, Alleyne M. 2022. Staying dry and cean: An insect’s guide to hydrophobicity. Insects 14:42. 10.3390/insects14010042

Benítez-Álvarez L, Tonzo V, Aristide L, Fernández R. 2026. Parallel genomic remodelling associated with independent terrestrialization events in arthropods. Mol. Ecol. 35:e70231. 10.1111/mec.70231

Bentley VL, Pérez-Moreno JL, Durica DS, Mykles DL. 2026. Characterization of the crustacean methyl farnesoate transcriptional signaling genes. Int. J. Mol. Sci. 27:1215. 10.3390/ijms27031215

Bernot JP, Avdeyev P, Zamyatin A, Dreyer N, Alexeev N, Pérez-Losada M, Crandall KA. 2022. Chromosome-level genome assembly, annotation, and phylogenomics of the gooseneck barnacle Pollicipes pollicipes. GigaScience 11:giac021. 10.1093/gigascience/giac021

Bernot JP, Owen CL, Wolfe JM, Meland K, Olesen J, Crandall KA. 2023. Major revisions in pancrustacean phylogeny and evidence of sensitivity to taxon sampling. Mol. Biol. Evol. 40:msad175. 10.1093/molbev/msad175

Betancourt JL, Rodríguez-Ramos T, Dixon B. 2024. Pattern recognition receptors in Crustacea: immunological roles under environmental stress. Front. Immunol. 15:1474512. 10.3389/fimmu.2024.1474512

Bishop CD, Erezyilmaz DF, Flatt T, Georgiou CD, Hadfield MG, Heyland A, Hodin J, Jacobs MW, Maslakova SA, Pires A, et al. 2006. What is metamorphosis? Integr. Comp. Biol. 46:655–661. 10.1093/icb/icl004

Blaxter M, Lewin HA, DiPalma F, Challis R, Da Silva M, Durbin R, Formenti G, Franz N, Guigo R, Harrison PW, et al. 2025. The Earth BioGenome Project Phase II: illuminating the eukaryotic tree of life. Front. Sci. 3:1514835. 10.3389/fsci.2025.1514835

Blum M, Andreeva A, Florentino LC, Chuguransky SR, Grego T, Hobbs E, Pinto BL, Orr A, Paysan-Lafosse T, Ponamareva I, et al. 2025. InterPro: the protein sequence classification resource in 2025. Nucleic Acids Res. 53:D444–D456. 10.1093/nar/gkae1082

Campli G, Volovych O, Kim K, Veldsman WP, Drage HB, Sheizaf I, Lynch S, Chipman AD, Daley AC, Robinson-Rechavi M, et al. 2024. The moulting arthropod: a complete genetic toolkit review. Biol. Rev. 99:2338–2375. 10.1111/brv.13123

Capella-Gutiérrez S, Silla-Martínez JM, Gabaldón T. 2009. trimAl: a tool for automated alignment trimming in large-scale phylogenetic analyses. Bioinforma. Oxf. Engl. 25:1972– 1973. 10.1093/bioinformatics/btp348

Chafino S, Giannios P, Casanova J, Martín D, Franch-Marro X. 2023. Antagonistic role of the BTB-zinc finger transcription factors Chinmo and Broad-Complex in the juvenile/pupal transition and in growth control. eLife 12:e84648. 10.7554/eLife.84648

Cheng S, Park Y, Kurleto JD, Jeon M, Zinn K, Thornton JW, Özkan E. 2019. Family of neural wiring receptors in bilaterians defined by phylogenetic, biochemical, and structural evidence. Proc. Natl. Acad. Sci. 116:9837–9842. 10.1073/pnas.1818631116

Chipman AD. 2020. The evolution of the gene regulatory networks patterning the Drosophila blastoderm. Curr. Top. Dev. Biol. 139:297–324. 10.1016/bs.ctdb.2020.02.004

Chipman AD. 2025. The development and evolution of arthropod tagmata. Proc. R. Soc. B Biol. Sci. 292:20242950. 10.1098/rspb.2024.2950

Chou A, Lin C, Cronin TW. 2020. Visual metamorphoses in insects and malacostracans: Transitions between an aquatic and terrestrial life. Arthropod Struct. Dev. 59:100974. 10.1016/j.asd.2020.100974

Clark E, Peel AD, Akam M. 2019. Arthropod segmentation. Development 146:dev170480. 10.1242/dev.170480

Culurgioni S, Alfieri A, Pendolino V, Laddomada F, Mapelli M. 2011. Inscuteable and NuMA proteins bind competitively to Leu-Gly-Asn repeat-enriched protein (LGN) during asymmetric cell divisions. Proc. Natl. Acad. Sci. U. S. A. 108:20998–21003. 10.1073/pnas.1113077108

Edgar RC. 2004. MUSCLE: multiple sequence alignment with high accuracy and high throughput. Nucleic Acids Res. 32:1792–1797. 10.1093/nar/gkh340

Fernandez-Nicolas A, Machaj G, Ventos-Alfonso A, Pagone V, Minemura T, Ohde T, Daimon T, Ylla G, Belles X. 2023. Reduction of embryonic E93 expression as a hypothetical driver of the evolution of insect metamorphosis. Proc. Natl. Acad. Sci. 120:e2216640120. 10.1073/pnas.2216640120

Feron R, Waterhouse RM. 2022. Assessing species coverage and assembly quality of rapidly accumulating sequenced genomes. GigaScience 11:giac006. 10.1093/gigascience/giac006

Gao Y, Zhang X, Wei J, Sun X, Yuan J, Li F, Xiang J. 2015. Whole transcriptome analysis provides insights into molecular mechanisms for molting in Litopenaeus vannamei. PLOS ONE 10:e0144350. 10.1371/journal.pone.0144350

Green J, Akam M. 2013. Evolution of the pair rule gene network: Insights from a centipede. Dev. Biol. 382:235–245. 10.1016/j.ydbio.2013.06.017

Haug JT. 2020a. Why the term “larva” is ambiguous, or what makes a larva? Acta Zool. 101:167–188. 10.1111/azo.12283

Haug JT. 2020b. Metamorphosis in crustaceans. In: Anger K, Harzsch S, Thiel M, editors. Developmental Biology and Larval Ecology. 1st ed. Oxford University Press. p. 255–284. 10.1093/oso/9780190648954.003.0009

Hoang DT, Chernomor O, von Haeseler A, Minh BQ, Vinh LS. 2018. UFBoot2: Improving the Ultrafast Bootstrap Approximation. Mol. Biol. Evol. 35:518–522. 10.1093/molbev/msx281

Hothorn T, Hornik K, Wiel MA van de, Zeileis A. 2008. Implementing a class of permutation tests: The coin package. J. Stat. Softw. 28:1–23. 10.18637/jss.v028.i08

Janssen R, Damen WGM, Budd GE. 2012. Expression of pair rule gene orthologs in the blastoderm of a myriapod: evidence for pair rule-like mechanisms? BMC Dev. Biol. 12:15. 10.1186/1471-213X-12-15

Jindra M. 2019. Where did the pupa come from? The timing of juvenile hormone signalling supports homology between stages of hemimetabolous and holometabolous insects. Philos. Trans. R. Soc. B Biol. Sci. 374:20190064. 10.1098/rstb.2019.0064

Kalyaanamoorthy S, Minh BQ, Wong TKF, von Haeseler A, Jermiin LS. 2017. ModelFinder: fast model selection for accurate phylogenetic estimates. Nat. Methods 14:587–589. 10.1038/nmeth.4285

Kamiyama T, Niwa R. 2022. Transcriptional regulators of ecdysteroid biosynthetic enzymes and their roles in insect development. Front. Physiol. 13. 10.3389/fphys.2022.823418

Kamsoi O, Ventos-Alfonso A, Casares F, Almudi I, Belles X. 2021. Regulation of metamorphosis in neopteran insects is conserved in the paleopteran Cloeon dipterum (Ephemeroptera). Proc. Natl. Acad. Sci. 118:e2105272118. 10.1073/pnas.2105272118

Kelly TR, Fitzgibbon QP, Smith GG, Banks TM, Ventura T. 2023. Tropical rock lobster (Panulirus ornatus) uses chemoreception via the antennular lateral flagellum to identify conspecific ecdysis. Sci. Rep. 13:12409. 10.1038/s41598-023-39567-8

Kim J-H, Kim Hyunkyong, Kim Heesoo, Chan BKK, Kang S, Kim W. 2019. Draft genome assembly of a fouling barnacle, Amphibalanus amphitrite (Darwin, 1854): The first reference genome for Thecostraca. Front. Ecol. Evol. 7. 10.3389/fevo.2019.00465

Kim K, Campli G, Joye S, Chipman AD, Waterhouse RM, Robinson-Rechavi M. 2026. Moulting in Pancrustacea is characterised by both deeply conserved and recently evolved gene modules. 10.64898/2026.01.15.699689

Kor G, Mengal K, Buřič M, Kozák P, Niksirat H. 2023. Comparative ultrastructure of the antennae and sensory hairs in six species of crayfish. PeerJ 11:e15006. 10.7717/peerj.15006

Krause C, Wolf C, Hemphälä J, Samakovlis C, Schuh R. 2006. Distinct functions of the leucine-rich repeat transmembrane proteins capricious and tartan in the Drosophila tracheal morphogenesis. Dev. Biol. 296:253–264. 10.1016/j.ydbio.2006.04.462

Kumar S, Suleski M, Craig JM, Kasprowicz AE, Sanderford M, Li M, Stecher G, Hedges SB. 2022. TimeTree 5: An Expanded Resource for Species Divergence Times. Mol. Biol. Evol. 39:msac174. 10.1093/molbev/msac174

Kuznetsov D, Tegenfeldt F, Manni M, Seppey M, Berkeley M, Kriventseva EV, Zdobnov EM. 2023. OrthoDB v11: annotation of orthologs in the widest sampling of organismal diversity. Nucleic Acids Res. 51:D445–D451. 10.1093/nar/gkac998

Liu L, Liu X, Fu Y, Fang W, Wang C. 2022. Whole-body transcriptome analysis provides insights into the cascade of sequential expression events involved in growth, immunity, and metabolism during the molting cycle in Scylla paramamosain. Sci. Rep. 12:11395. 10.1038/s41598-022-14783-w

MacLaren CM, Evans TA, Alvarado D, Duffy JB. 2004. Comparative analysis of the Kekkon molecules, related members of the LIG superfamily. Dev. Genes Evol. 214:360–366. 10.1007/s00427-004-0414-4

Manni M, Berkeley MR, Seppey M, Zdobnov EM. 2021. BUSCO: Assessing genomic data quality and beyond. Curr. Protoc. 1:e323. 10.1002/cpz1.323

Martin JW, Olesen J, Høeg JT. 2014. Atlas of crustacean larvae. Johns Hopkins University Press 10.1353/book.31448

Mazzoni CJ, Ciofi C, Waterhouse RM. 2023. Biodiversity: an atlas of European reference genomes. Nature 619:252–252. 10.1038/d41586-023-02229-w

Mellon D. 2012. Smelling, feeling, tasting and touching: behavioral and neural integration of antennular chemosensory and mechanosensory inputs in the crayfish. J. Exp. Biol. 215:2163–2172. 10.1242/jeb.069492

Mendes FK, Vanderpool D, Fulton B, Hahn MW. 2021. CAFE 5 models variation in evolutionary rates among gene families. Bioinformatics 36:5516–5518. 10.1093/bioinformatics/btaa1022

Milán M, Weihe U, Pérez L, Cohen SM. 2001. The LRR proteins Capricious and Tartan mediate cell interactions during DV boundary formation in the Drosophila wing. Cell 106:785–794. 10.1016/S0092-8674(01)00489-5

Minh BQ, Schmidt HA, Chernomor O, Schrempf D, Woodhams MD, von Haeseler A, Lanfear R. 2020. IQ-TREE 2: New models and efficient methods for phylogenetic inference in the genomic era. Mol. Biol. Evol. 37:1530–1534. 10.1093/molbev/msaa015

Moran NA. 1994. Adaptation and constraint in the complex life cycles of animals. Annu. Rev. Ecol. Syst. 25:573–600.

Myasnikova E, Spirov A. 2020. Gene regulatory networks in Drosophila early embryonic development as a model for the study of the temporal identity of neuroblasts. Biosystems 197:104192. 10.1016/j.biosystems.2020.104192

Naggan Perl T, Schmid BGM, Schwirz J, Chipman AD. 2013. The evolution of the knirps family of transcription factors in arthropods. Mol. Biol. Evol. 30:1348–1357. 10.1093/molbev/mst046

Nguyen L-T, Schmidt HA, von Haeseler A, Minh BQ. 2015. IQ-TREE: A Fast and Effective Stochastic Algorithm for Estimating Maximum-Likelihood Phylogenies. Mol. Biol. Evol. 32:268–274. 10.1093/molbev/msu300

Nolan RB, Fan J-Y, Price JL. 2024. Circadian rhythms in the Drosophila eye may regulate adaptation of vision to light intensity. Front. Neurosci. 18. 10.3389/fnins.2024.1401721

Nunes C, Sucena É, Koyama T. 2021. Endocrine regulation of immunity in insects. FEBS J. 288:3928–3947. 10.1111/febs.15581

Okamoto N, Yamanaka N. 2020. Steroid hormone entry into the brain requires a membrane transporter in Drosophila. Curr. Biol. CB 30:359–366.e3. 10.1016/j.cub.2019.11.085

Okude G, Moriyama M, Kawahara-Miki R, Yajima S, Fukatsu T, Futahashi R. 2022. Molecular mechanisms underlying metamorphosis in the most-ancestral winged insect. Proc. Natl. Acad. Sci. 119:e2114773119. 10.1073/pnas.2114773119

Olesen J. 2018. Crustacean life cycles—Developmental strategies and environmental adaptations. In: Thiel M, Wellborn GA, editors. Life Histories. 1st ed. Oxford University PressNew York. p. 1–34. 10.1093/oso/9780190620271.003.0001

Öztürk-Çolak A, Marygold SJ, Antonazzo G, Attrill H, Goutte-Gattat D, Jenkins VK, Matthews BB, Millburn G, Dos Santos G, Tabone CJ, et al. 2024. FlyBase: updates to the Drosophila genes and genomes database. Wood V, editor. GENETICS 227:iyad211. 10.1093/genetics/iyad211

Paradis E, Schliep K. 2019. ape 5.0: an environment for modern phylogenetics and evolutionary analyses in R. Bioinforma. Oxf. Engl. 35:526–528. 10.1093/bioinformatics/bty633

Reding K, Chen M, Lu Y, Cheatle Jarvela AM, Pick L. 2019. Shifting roles of Drosophila pair-rule gene orthologs: segmental expression and function in the milkweed bug Oncopeltus fasciatus. Development 146:dev181453. 10.1242/dev.181453

Reijnders MJMF, Waterhouse RM. 2021. Summary visualizations of gene ontology terms with GO-Figure! *Front*. Bioinforma. 1. 10.3389/fbinf.2021.638255

Roer R, Abehsera S, Sagi A. 2015. Exoskeletons across the Pancrustacea: Comparative morphology, physiology, biochemistry and genetics. Integr. Comp. Biol. 55:771–791. 10.1093/icb/icv080

San-Juán BP, Baonza A. 2011. The bHLH factor deadpan is a direct target of Notch signaling and regulates neuroblast self-renewal in Drosophila. Dev. Biol. 352:70–82. 10.1016/j.ydbio.2011.01.019

Sato M, Yasugi T, Trush O. 2019. Temporal patterning of neurogenesis and neural wiring in the fly visual system. Neurosci. Res. 138:49–58. 10.1016/j.neures.2018.09.009

Seppey M, Ioannidis P, Emerson BC, Pitteloud C, Robinson-Rechavi M, Roux J, Escalona HE, McKenna DD, Misof B, Shin S, et al. 2019. Genomic signatures accompanying the dietary shift to phytophagy in polyphagan beetles. Genome Biol. 20:98. 10.1186/s13059-019-1704-5

Shieh B-H, Kristaponyte I, Hong Y. 2014. Distinct roles of arrestin 1 protein in photoreceptors during Drosophila development. J. Biol. Chem. 289:18526–18534. 10.1074/jbc.M114.571224

Shimozono M, Osaka J, Kato Y, Araki T, Kawamura H, Takechi H, Hakeda-Suzuki S, Suzuki T. 2019. Cell surface molecule, Klingon, mediates the refinement of synaptic specificity in the Drosophila visual system. Genes Cells Devoted Mol. Cell. Mech. 24:496–510. 10.1111/gtc.12703

Snodgrass RE. 1956. Crustacean metamorphoses. Smithson. Misc. Collect. 131:1–78.

Spitzner F, Meth R, Krüger C, Nischik E, Eiler S, Sombke A, Torres G, Harzsch S. 2018. An atlas of larval organogenesis in the European shore crab Carcinus maenas L. (Decapoda, Brachyura, Portunidae). Front. Zool. 15:27. 10.1186/s12983-018-0271-z

Sun DA, Patel NH. 2019. The amphipod crustacean Parhyale hawaiensis: An emerging comparative model of arthropod development, evolution, and regeneration. *WIREs Dev*. Biol. 8:e355. 10.1002/wdev.355

Ten Brink H, De Roos AM, Dieckmann U. 2019. The evolutionary ecology of metamorphosis. Am. Nat. 193:E116–E131. 10.1086/701779

Thomas GWC, Dohmen E, Hughes DST, Murali SC, Poelchau M, Glastad K, Anstead CA, Ayoub NA, Batterham P, Bellair M, et al. 2020. Gene content evolution in the arthropods. Genome Biol. 21:15. 10.1186/s13059-019-1925-7

True JR, Carroll SB. 2002. Gene co-option in physiological and morphological evolution. Annu. Rev. Cell Dev. Biol. 18:53–80. 10.1146/annurev.cellbio.18.020402.140619

Truman JW. 2019. The evolution of insect metamorphosis. Curr. Biol. 29:R1252–R1268. 10.1016/j.cub.2019.10.009

Truman JW, Price J, Miyares RL, Lee T. 2023. Metamorphosis of memory circuits in Drosophila reveals a strategy for evolving a larval brain. eLife 12:e80594. 10.7554/eLife.80594

Ulian-Benitez S, Bishop S, Foldi I, Wentzell J, Okenwa C, Forero MG, Zhu B, Moreira M, Phizacklea M, McIlroy G, et al. 2017. Kek-6: A truncated-Trk-like receptor for Drosophila neurotrophin 2 regulates structural synaptic plasticity. PLoS Genet. 13:e1006968. 10.1371/journal.pgen.1006968

Urbach R, Jussen D, Technau GM. 2016. Gene expression profiles uncover individual identities of gnathal neuroblasts and serial homologies in the embryonic CNS of Drosophila. Development 143:1290–1301. 10.1242/dev.133546

Vea IM, Minakuchi C. 2021. Atypical insects: molecular mechanisms of unusual life history strategies. Curr. Opin. Insect Sci. 43:46–53. 10.1016/j.cois.2020.09.016

Ventura T, Manor R, Aflalo ED, Chalifa-Caspi V, Weil S, Sharabi O, Sagi A. 2013. Post-embryonic transcriptomes of the prawn Macrobrachium rosenbergii: Multigenic succession through metamorphosis. PLoS ONE 8:e55322. 10.1371/journal.pone.0055322

Wang J, Dahmann C. 2020. Establishing compartment boundaries in Drosophila wing imaginal discs: An interplay between selector genes, signaling pathways and cell mechanics. Semin. Cell Dev. Biol. 107:161–169. 10.1016/j.semcdb.2020.07.008

Waterhouse RM, Tegenfeldt F, Li J, Zdobnov EM, Kriventseva EV. 2013. OrthoDB: a hierarchical catalog of animal, fungal and bacterial orthologs. Nucleic Acids Res. 41:D358–D365. 10.1093/nar/gks1116

Wei J, Pisani D, Donoghue PCJ, Álvarez-Presas M, Paps J. 2026. Convergent genome evolution shaped the emergence of terrestrial animals. Nature 649:638–646. 10.1038/s41586-025-09722-4

Wexler J, Pick L, Chipman A. 2023. Segmental expression of two ecdysone pathway genes during embryogenesis of hemimetabolous insects. Dev. Biol. 498:87–96. 10.1016/j.ydbio.2023.03.008

Wong TKF, Kalyaanamoorthy S, Meusemann K, Yeates DK, Misof B, Jermiin LS. 2020. A minimum reporting standard for multiple sequence alignments. NAR Genomics Bioinforma. 2: lqaa024. 10.1093/nargab/lqaa024

Yamasaki Y, Lim Y-M, Niwa N, Hayashi S, Tsuda L. 2011. Robust specification of sensory neurons by dual functions of charlatan, a Drosophila NRSF/REST-like repressor of extramacrochaetae and hairy. Genes Cells Devoted Mol. Cell. Mech. 16:896–909. 10.1111/j.1365-2443.2011.01537.x

Ylla G, Piulachs M-D, Belles X. 2018. Comparative transcriptomics in two extreme neopterans reveals general trends in the evolution of modern insects. iScience 4:164–179. 10.1016/j.isci.2018.05.017

Yuan J, Zhang Xiaojun, Zhang Xiaoxi, Sun Y, Liu C, Li S, Yu Y, Zhang C, Jin S, Wang M, et al. 2024. An ancient whole-genome duplication in barnacles contributes to their diversification and intertidal sessile life adaptation. J. Adv. Res. 62:91–103. 10.1016/j.jare.2023.09.015

