## Supplementary Figures for "Convergent gene family evolution underpins repeated transitions to metamorphic development across Pancrustacea"

**Supplementary Figure 1: Species tree and orthology**

**(A)** The species phylogeny shows the relationships and estimated divergence times of the complete dataset of 54 arthropod species. The tree is annotated with the 26 represented orders. The phylogeny was estimated using a super-alignment of protein sequences from 121 single-copy orthologues and time-calibrated with ten calibration nodes. The Crustacea (yellow) and Hexapoda (green) are highlighted, where hexapods and the paraphyletic crustaceans together form the monophyletic Pancrustacea. Orders are annotated according to their developmental mode: metamorphic lineages (red), non-metamorphic sister lineages (blue). Note that here Podocopida is not, in the strict sense, a sister lineage to Thecostraca, but rather an outgroup lineage for which available data mean it can be used for the phylogenomic contrasts. **(B)** Orthology delineation at the level of the last common ancestor (LCA) of the complete dataset of 54 arthropod species resulted in the orthologous group (OG) dataset comprising a total of 42'841 OGs and 782'987 genes. The bars show the counts of genes in OGs classified as single-copy (blue) or multi-copy (green, yellow) orthologues, or for which no orthologues could be identified (grey).

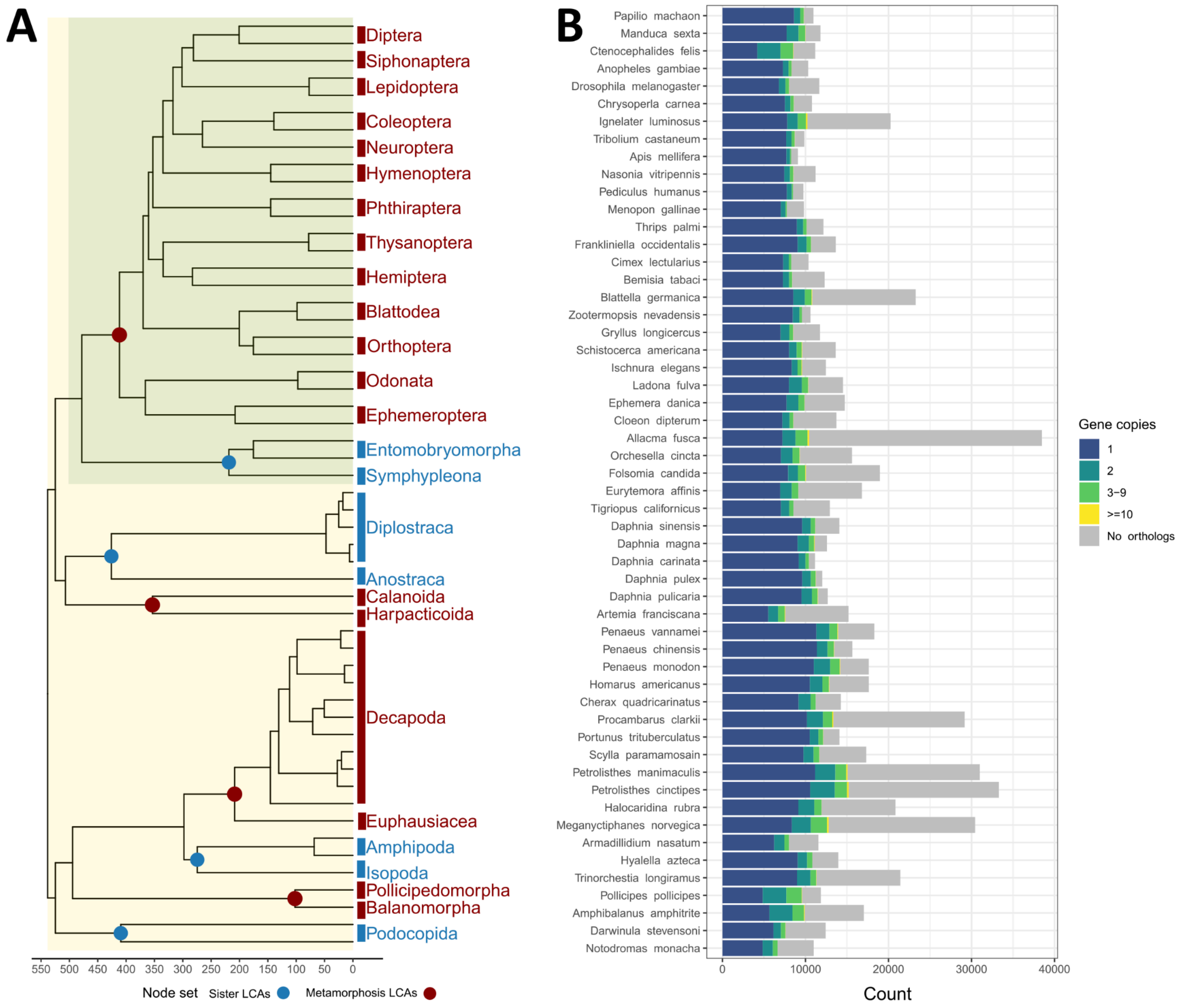

**Supplementary Figure 2: Branch lengths of metamorphosis LCAs and control nodes**

**(A)** Branch lengths expressed as millions of years, for each node of the node sets (deep nodes in grey, metamorphosis LCAs in red, and sister LCAs in blue), separating them from their closest ancestor nodes. All pairs are not significant according to Wilcoxon-Mann-Whitney Tests with Permutation (Metamorphosis LCAs - Sister LCAs p-value = 0.56, Metamorphosis LCAs - Deep nodes p-value = 0.06, Sister LCAs - Deep nodes p-value = 0.14). The boxplots show the median, first and third quartiles, and lower and upper extremes of the distribution (1.5 × Interquartile range). **(B)** Branch lengths expressed as millions of years, summed over the full path from the tip to each node of the node sets representing their age of appearance. Asterisks indicate Wilcoxon-Mann-Whitney Tests with Permutation, *p<0.05 (Metamorphosis LCAs - Sister LCAs p-value = 0.39, Metamorphosis LCAs - Deep nodes p-value = 0.03, Sister LCAs - Deep nodes p-value = 0.03). The boxplots show the median, first and third quartiles, and lower and upper extremes of the distribution (1.5 × Interquartile range).

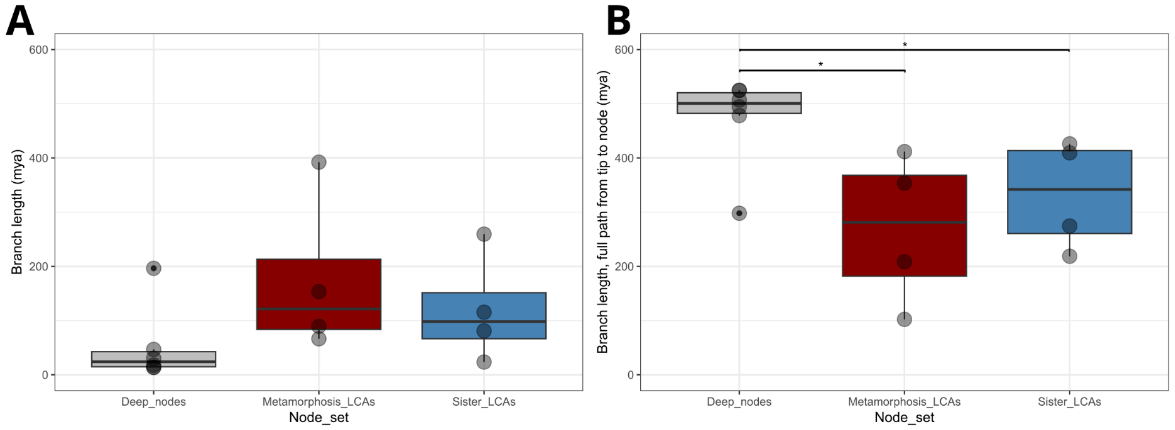

**Supplementary Figure 3: Counts of gene family evolutionary events at nodes of interest.**

**A)** Partitioning of the orthologous groups (OGs) by phyletic age, defined as the last common ancestor (LCA) of all the species included in the OG. Ages include the Pancrustacea ancestor, the LCAs of metamorphosis lineages (Metamorphosis LCAs, n=4), their sister lineages (Sister LCAs, n=4), their ancestors speciating since the Pancrustacea ancestor (Deep nodes, n=6), and all the nodes descendant from the Metamorphosis and Sister LCAs (Younger nodes, n=38). **(B)** Counts of ancient and widespread OGs (present in the Pancrustacea LCA and with orthologues in at least 85% of the species) that experienced expansions (gene gains) or contractions (gene losses) at the Metamorphosis LCAs, Sister LCAs, Deep nodes, and Younger nodes.

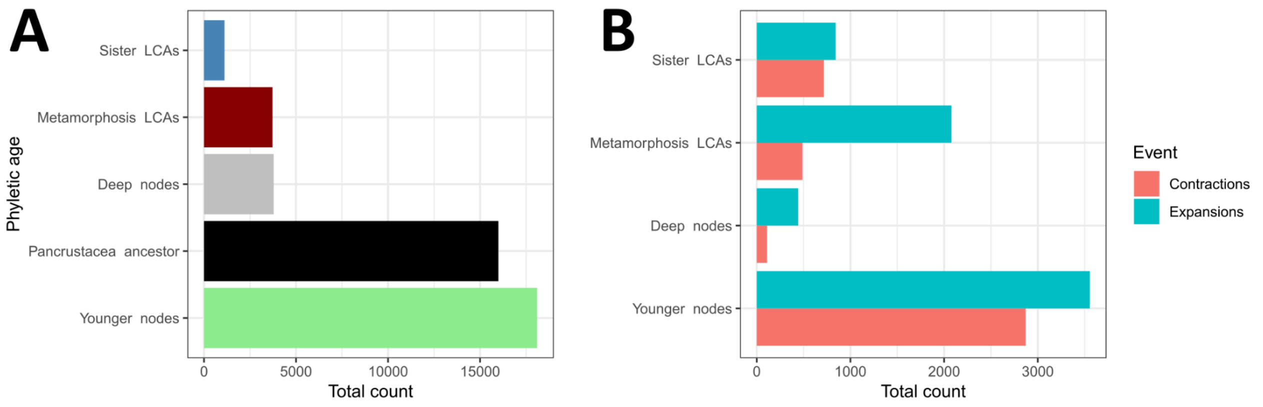

**Supplementary Figure 4: Distribution of evolutionary rates of gene families dated at the Pancrustacea LCA**

Evolutionary rates quantify sequence-level dynamics in terms of the levels of protein sequence divergence amongst orthologous group (OG) member genes. OG evolutionary rates were computed using OrthoLoger ancillary tools: pairwise protein alignments are used to calculate average of inter-species identities for each OG, normalised over the average identity of all inter-species BRH, as defined by OrthoDB. The violin plot shows the smoothed kernel density of the distribution, and the boxplot within shows the median as well as the first and third quartiles of the distribution.

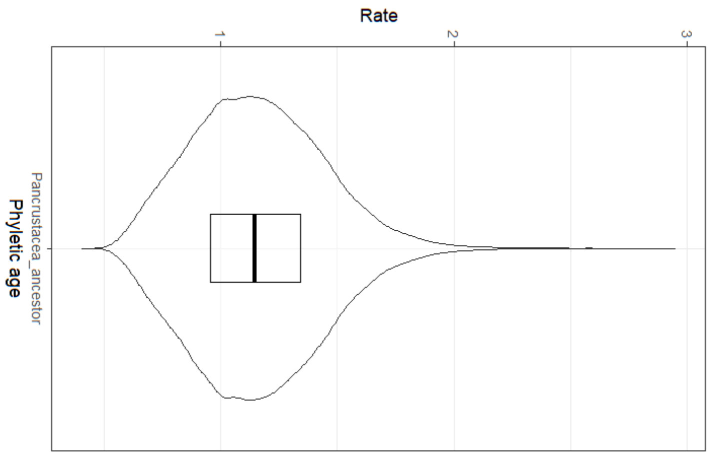

**Supplementary Figure 5: Semantic similarity plot summarising biological functions enriched amongst gene families emerging at the Metamorphosis LCAs.**

Summary visualisations of lists of enriched Gene Ontology (GO) terms associated with orthologous groups (OGs) emerging at Metamorphosis LCAs, contrasted with OGs emerging at any Sister LCA and any offspring node of Metamorphosis and Sister LCAs (Younger nodes). It should be noted that amongst the full set of emerging OGs (i.e. gene family births, gene families having the phyletic age of a node in the respective set) there were generally low proportions annotated with GO terms: Metamorphosis LCAs 48% Insecta, 44% Eucarida, 40% Copepoda, 27% Thecostraca, and Sister LCAs 46% non-Insecta hexapods, 26% Peracarida, 58% Branchiopoda, and 33% Ostracoda. Therefore, the contrasts of gene families that emerged at any of the Metamorphosis LCAs or of any of the Sister LCAs with births at other internal nodes of the phylogeny include many OGs with no functional annotations. Larger circles group a higher number of closely related terms together in the same semantic space, the smallest circles representing a single term each. The positioning of the circles along the X and Y axes maximises functional distinctions in the semantic space, where more similar functions are closer together on the plot. Circle colours represent significance values (log10 p-value) from the GO enrichment analysis, according to the legend. Number labels on each circle correspond to the presented list of GO term names, ordered by significance values.

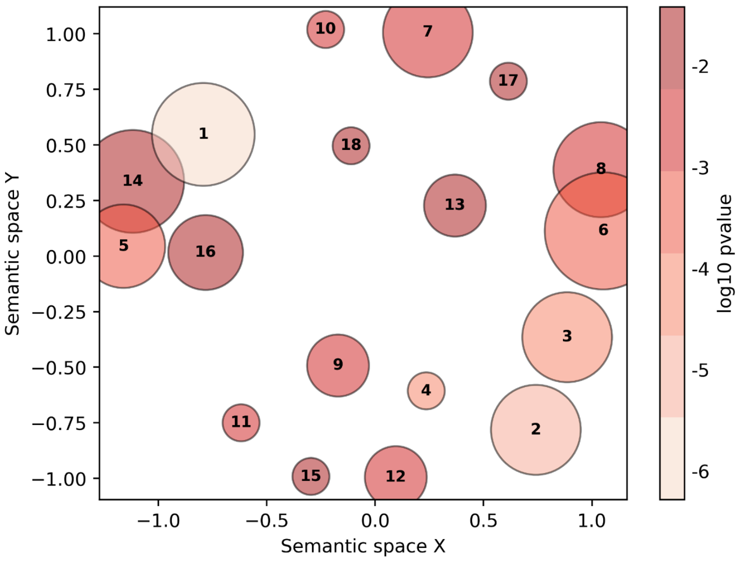

1. Wnt signaling pathway
2. response to organic cyclic compound
3. sodium ion transport
4. protein peptidyl-prolyl isomerization
5. positive regulation of innate immune response
6. cellular component assembly involved in morphogenesis
7. metamorphosis
8. chitin-based cuticle development
9. DNA transposition
10. circadian rhythm
11. organic hydroxy compound metabolic process
12. nucleobase-containing compound catabolic process
13. developmental growth involved in morphogenesis
14. positive regulation of cell population proliferation
15. carbohydrate biosynthetic process
16. regulation of anatomical structure morphogenesis
17. intracellular chemical homeostasis
18. establishment or maintenance of cell polarity

**Supplementary Figure 6: Common and unique OGs expanding at Metamorphosis LCAs.**

The upset plot shows the partitioning of the 856 orthologous groups (OGs) annotated with the 60 Gene Ontology (GO) terms found to be enriched amongst the sets of expanding OGs at all four Metamorphosis LCAs according to which node or nodes the expansion events were inferred to have taken place.

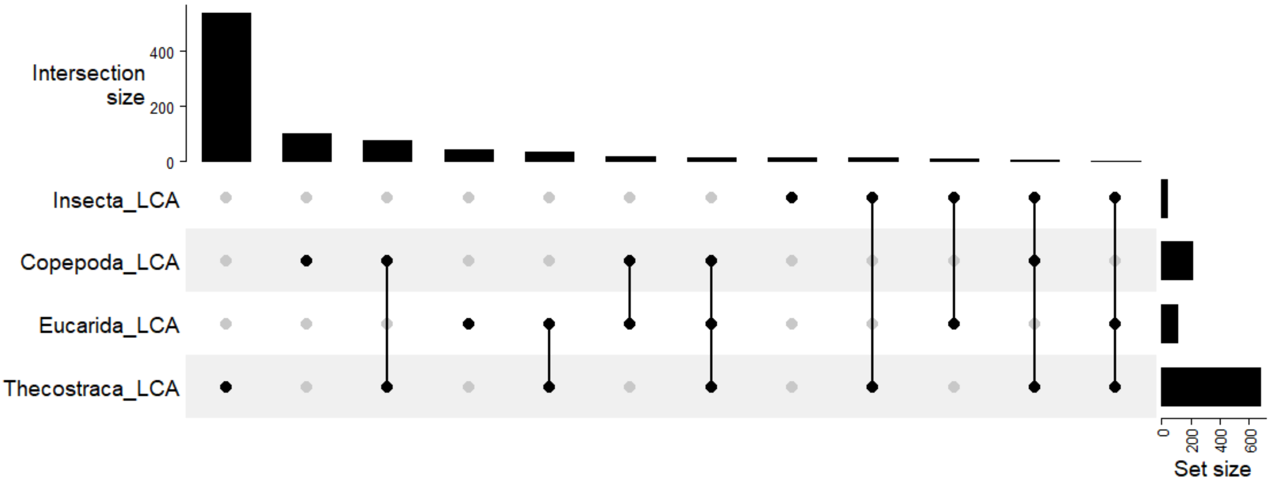
