## Supplementary Tables for "Convergent gene family evolution underpins repeated transitions to metamorphic development across Pancrustacea"

| Assembly Accession | TaxId | Species | Arthropoda Complete BUSCO | Arthropoda Single BUSCO | Arthropoda Duplicated BUSCO | Number of protein coding genes | Arthropoda Complete BUSCO Proteomes | Arthropoda Single BUSCO Proteomes | Arthropoda Duplicated BUSCO Proteomes |
|---|---|---|---|---|---|---|---|---|---|
| GCA_000376725.2 | 123851 | Ladona fulva | 93.7 | 92.3 | 1.4 | 18816 | 78.2 | 76.6 | 1.6 |
| GCA_000507165.2 | 1049336 | Ephemera danica | 94.5 | 91.6 | 2.9 | 18562 | 85.8 | 84.3 | 1.5 |
| GCA_001718145.1 | 48709 | Orchesella cincta | 93 | 88.4 | 4.6 | 20247 | 90 | 85.2 | 4.8 |
| GCA_003018175.1 | 6973 | Blattella germanica | 96.5 | 94 | 2.5 | 28676 | 84.9 | 82.9 | 2 |
| GCA_006783055.1 | 1923959 | Trinorchestia longiramus | 86.4 | 83.1 | 3.3 | 26080 | 86.5 | 83 | 3.5 |
| GCA_007210705.1 | 6832 | Tigriopus californicus | 93.5 | 92.8 | 0.7 | 15577 | 91.9 | 90.6 | 1.3 |
| GCA_009176605.1 | 96803 | Armadillidium nasatum | 89.3 | 83.1 | 6.2 | 14636 | 75.3 | 70.5 | 4.8 |
| GCA_009805615.1 | 1232801 | Amphibalanus amphitrite | 92.5 | 64 | 28.5 | 25581 | 85.2 | 61.3 | 23.9 |
| GCA_011009095.1 | 2038154 | Ignelater luminosus | 96.9 | 94.4 | 2.5 | 27558 | 94.5 | 92.1 | 2.4 |
| GCA_013167095.2 | 1820382 | Daphnia sinensis | 98.5 | 96.3 | 2.2 | 16955 | 97.4 | 94.8 | 2.6 |
| GCA_032884065.1 | 6661 | Artemia franciscana | 89.4 | 81.5 | 7.9 | 20122 | 81.3 | 75.4 | 5.9 |
| GCA_033782935.1 | 88211 | Petrolisthes cinctipes | 92 | 75.2 | 16.8 | 44511 | 94.4 | 77 | 17.4 |
| GCA_034508575.1 | 1843537 | Petrolisthes manimaculis | 92.4 | 79.7 | 12.7 | 40296 | 95.1 | 82.1 | 13 |
| GCA_037179515.1 | 373956 | Halocaridina rubra | 88.5 | 86.2 | 2.3 | 25341 | 82.5 | 81.4 | 1.1 |
| GCA_038098605.1 | 2509291 | Gryllus longicercus | 99 | 90.3 | 8.7 | 14730 | 95.6 | 87.6 | 8 |
| GCA_038387795.1 | 85552 | Scylla paramamosain | 95.1 | 93.5 | 1.6 | 20946 | 89.9 | 88.4 | 1.5 |
| GCA_038502225.1 | 27406 | Cherax quadricarinatus | 88.8 | 86.9 | 1.9 | 17532 | 88.3 | 86.4 | 1.9 |
| GCA_040285465.1 | 328185 | Menopon gallinae | 98.8 | 98.2 | 0.6 | 10891 | 88.5 | 87.5 | 1 |
| GCA_902829235.1 | 197152 | Cloeon dipterum | 97.8 | 95.4 | 2.4 | 16299 | 97 | 94.5 | 2.5 |
| GCA_905338385.1 | 69355 | Darwinula stevensoni | 90.3 | 88.1 | 2.2 | 15334 | 84.5 | 82.5 | 2 |
| GCA_905338405.1 | 399045 | Notodromas monacha | 86.2 | 81.1 | 5.1 | 13723 | 80.8 | 76.7 | 4.1 |
| GCA_910591605.1 | 39272 | Allacma fusca | 91 | 89.2 | 1.8 | 47610 | 91.8 | 89.7 | 2.1 |
| GCA_918797505.1 | 7038 | Bemisia tabaci | 98.5 | 96.1 | 2.4 | 14348 | 96.3 | 94.1 | 2.2 |
| GCA_964058975.1 | 48144 | Meganyctiphanes norvegica | 75.7 | 71.7 | 4 | 42189 | 85.3 | 82 | 3.3 |
| GCF_000001215.4 | 7227 | Drosophila melanogaster | 99.6 | 99.1 | 0.5 | 13986 | 99.9 | 99 | 0.9 |
| GCF_000006295.1 | 121224 | Pediculus humanus | 97.1 | 96.9 | 0.2 | 10758 | 97 | 96.7 | 0.3 |
| GCF_000591075.1 | 88015 | Eurytemora affinis | 90.9 | 88.6 | 2.3 | 20716 | 93.8 | 91.7 | 2.1 |
| GCF_000648675.2 | 79782 | Cimex lectularius | 99.4 | 97.8 | 1.6 | 11949 | 99.5 | 98.2 | 1.3 |
| GCF_000696155.1 | 136037 | Zootermopsis nevadensis | 98.9 | 98.5 | 0.4 | 12381 | 98.4 | 98.1 | 0.3 |
| GCF_000697945.3 | 133901 | Frankliniella occidentalis | 98.3 | 97.3 | 1 | 16516 | 99.5 | 97.9 | 1.6 |
| GCF_000764305.2 | 294128 | Hyalella azteca | 93.7 | 93 | 0.7 | 17163 | 95.6 | 94.6 | 1 |
| GCF_002217175.1 | 158441 | Folsomia candida | 97.5 | 95.9 | 1.6 | 24221 | 97.9 | 96.1 | 1.8 |
| GCF_003254395.2 | 7460 | Apis mellifera | 99.8 | 99.8 | 0 | 9935 | 99.2 | 99 | 0.2 |
| GCF_003426905.1 | 7515 | Ctenocephalides felis | 96 | 69.4 | 26.6 | 18878 | 95.8 | 58.8 | 37 |
| GCF_009193385.2 | 7425 | Nasonia vitripennis | 98.8 | 97.1 | 1.7 | 13602 | 98.4 | 96.6 | 1.8 |
| GCF_011947565.2 | 41117 | Pollicipes pollicipes | 90.9 | 69.1 | 21.8 | 20444 | 93.3 | 57.3 | 36 |
| GCF_012932325.1 | 161013 | Thrips palmi | 96.9 | 95.9 | 1 | 14332 | 97.5 | 96.4 | 1.1 |
| GCF_014839805.1 | 7130 | Manduca sexta | 98.3 | 89.8 | 8.5 | 15967 | 99.3 | 89.1 | 10.2 |
| GCF_015228065.1 | 6687 | Penaeus monodon | 83.7 | 81.6 | 2.1 | 24092 | 91.3 | 87.3 | 4 |
| GCF_017591435.1 | 210409 | Portunus trituberculatus | 93.5 | 92.9 | 0.6 | 17292 | 97.2 | 96.3 | 0.9 |
| GCF_018991925.1 | 6706 | Homarus americanus | 93.2 | 92.2 | 1 | 22368 | 97.6 | 96.3 | 1.3 |
| GCF_019202785.1 | 139456 | Penaeus chinensis | 90.8 | 89.8 | 1 | 20076 | 95.8 | 94.5 | 1.3 |
| GCF_020631705.1 | 35525 | Daphnia magna | 98.6 | 94.4 | 4.2 | 16891 | 97.8 | 91.7 | 6.1 |
| GCF_021134715.1 | 6669 | Daphnia pulex | 98.1 | 97.9 | 0.2 | 15295 | 97.3 | 96.3 | 1 |
| GCF_021234035.1 | 35523 | Daphnia pulicaria | 98.8 | 97.2 | 1.6 | 16865 | 97.8 | 95.2 | 2.6 |
| GCF_021461395.2 | 7009 | Schistocerca americana | 98.2 | 96.2 | 2 | 17662 | 99.1 | 96.9 | 2.2 |
| GCF_022539665.1 | 120202 | Daphnia carinata | 98.7 | 97.6 | 1.1 | 13404 | 97.9 | 96.2 | 1.7 |
| GCF_031307605.1 | 7070 | Tribolium castaneum | 99.5 | 99.1 | 0.4 | 12172 | 99.3 | 98.5 | 0.8 |
| GCF_040958095.1 | 6728 | Procambarus clarkii | 94.4 | 93.2 | 1.2 | 37059 | 98.6 | 90.8 | 7.8 |
| GCF_042767895.1 | 6689 | Penaeus vannamei | 85 | 78.1 | 6.9 | 24066 | 92.1 | 88.8 | 3.3 |
| GCF_905475395.1 | 189513 | Chrysoperla carnea | 98 | 97.4 | 0.6 | 12985 | 98 | 97 | 1 |
| GCF_912999745.1 | 76193 | Papilio machaon | 99 | 98.1 | 0.9 | 13379 | 99.8 | 98.8 | 1 |
| GCF_921293095.1 | 197161 | Ischnura elegans | 99.2 | 98.1 | 1.1 | 15994 | 99.1 | 98.4 | 0.7 |
| GCF_943734735.2 | 180454 | Anopheles gambiae | 98.1 | 96.4 | 1.7 | 12519 | 99.3 | 98.2 | 1.1 |

**Supplementary Table 1: Genomic resources used to build the phylogenomics dataset.**

Assembly accessions, protein-coding gene counts, and Benchmarking Universal Single-Copy Orthologue (BUSCO) assessments from the publicly available annotated proteomes from the United States National Center for Biotechnology Information (NCBI).

| | |
|---|---|
| Number of sequences in the alignment | 54 |
| Number of sites in the alignment | 51083 |
| Pairs of sequences | 1431 |
| Completeness (C) score for the alignment (Ca) | 0.899276 |
| Maximum C-score for individual sequences (Cr max) | 0.972809 |
| Minimum C-score for individual sequences (Cr min) | 0.53509 |
| Maximum C-score for individual sites (Cc max) | 1 |
| Minimum C-score for individual sites (Cc min) | 0.018519 |
| Maximum C-score for pairs of sequences (Cij max, i!=j) | 0.959399 |
| Minimum C-score for pairs of sequences (Cij min, i!=j) | 0.347944 |

**Supplementary Table 2: Metrics of the concatenated super-alignment used for species phylogeny reconstruction.**

The species phylogeny was computed from 121 single-copy orthologues found in at least 95% of the total set of 54 species, detected by the BUSCO assessments performed for the A3Cat on the genome assemblies using the arthropoda_odb10 lineage dataset. A concatenated super-alignment was built from the trimmed individual multiple alignments (51’083 columns, 47’966 distinct patterns, 36’774 parsimony-informative, 6’920 singleton sites, 7’388 constant sites).

| Last Common Ancestor | Min estimate | Max estimate |
|---|---|---|
| Hexapoda | 425.4 | 478.1 |
| Insecta | 411.7 | 442.7 |
| Malacostraca | 254.2 | 444 |
| Branchiopoda | 365.1 | 491.7 |
| Hymenoptera,Lepidoptera | 310.7 | 389.7 |
| Root | 501.8 | 580 |

**Supplementary Table 3: Time estimates for species phylogeny tree calibration.**

Divergence time estimates were sourced from the TimeTree v5 database (Kumar et al. 2022) to time-calibrate the molecular species tree.

| Order | Class | Type of development | Transition Stage | Instars before transition | Characters at transition |
|---|---|---|---|---|---|
| Podocopida | Ostracoda | Partially anamorphic | none | | |
| Anostraca | Branchiopoda | Anamorphosis | none | | |
| Diplostraca | Branchiopoda | Direct | none | | |
| Isopoda | Malacostraca | Direct | none | | |
| Amphipoda | Malacostraca | Direct | none | | |
| Euphausiacea | Malacostraca | Metamorphic | furcilia | up to 3 | thoracopodal swimming |
| Decapoda | Malacostraca | Metamorphic | decapodid | up to 6 | thoracopodal swimming |
| Balanomorpha | Thecostraca | Metamorphic | cyprid | 6 | post-cephalic appendages for attachment phase |
| Pollicipedomorpha | Thecostraca | Metamorphic | cyprid | 6 | post-cephalic appendages for attachment phase |
| Calanoida | Copepoda | Metamorphic | copepodid | typically 6 | loss of NP on A2 coxa; naupliar somites separated |
| Harpacticoida | Copepoda | Metamorphic | copepodid | typically 7 | loss of NP on A2 coxa; naupliar somites separated |

**Supplementary Table 4: Development annotation in Crustacea based on Atlas of crustacean larvae *(*Martin, Olesen & Høeg 2014).**

Taxonomic ranking differences with NCBI taxonomy are reported. In the NCBI taxonomy, Balanomorpha is an order from subclass Cirripedia and class Thecostraca, while in Martin, Olesen & Høeg 2014 is a suborder from order Sessilia, from subclass Cirripedia and class Thecostraca. In the NCBI taxonomy, Pollicipedormorpha is an order. It has been recently created and includes species from the Pollicipes genus and the Anelasma, Capitulum, and Lithotrya genera, as shown in Bernot et al., 2022. In the NCBI taxonomy, Copepoda is a subclass from class Hexanauplia, while in Martin, Olesen & Høeg 2014 it is a class.

| Cluster representative | Cluster member | Cluster member description | Member P-value | Member IC | Member frequency |
|---|---|---|---|---|---|
| GO:0007186 | GO:0007186 | G protein-coupled receptor signaling pathway | 5.20E-07 | 2.32E+00 | 4.75E-03 |
| GO:0007186 | GO:0016055 | Wnt signaling pathway | 1.80E-02 | 3.17E+00 | 6.79E-04 |
| GO:0007186 | GO:0060070 | canonical Wnt signaling pathway | 3.95E-02 | 4.36E+00 | 4.33E-05 |
| GO:0007186 | GO:0051253 | negative regulation of RNA metabolic process | 2.81E-02 | 2.47E+00 | 3.39E-03 |
| GO:0007186 | GO:0048523 | negative regulation of cellular process | 1.64E-02 | 2.13E+00 | 7.35E-03 |
| GO:0007186 | GO:0010629 | negative regulation of gene expression | 2.82E-02 | 2.30E+00 | 5.04E-03 |
| GO:0007186 | GO:0043085 | positive regulation of catalytic activity | 2.61E-02 | 3.26E+00 | 5.44E-04 |
| GO:0007186 | GO:0065008 | regulation of biological quality | 4.34E-02 | 2.07E+00 | 8.58E-03 |
| GO:0014070 | GO:0014070 | response to organic cyclic compound | 2.10E-05 | 3.80E+00 | 1.57E-04 |
| GO:0014070 | GO:0071310 | cellular response to organic substance | 4.96E-03 | 3.49E+00 | 3.21E-04 |
| GO:0014070 | GO:0097305 | response to alcohol | 1.70E-04 | 4.25E+00 | 5.57E-05 |
| GO:0014070 | GO:0006281 | DNA repair | 1.18E-02 | 1.73E+00 | 1.85E-02 |
| GO:0014070 | GO:0006260 | DNA replication | 1.64E-02 | 2.06E+00 | 8.70E-03 |
| GO:0006814 | GO:0006814 | sodium ion transport | 4.30E-05 | 2.58E+00 | 2.63E-03 |
| GO:0006814 | GO:0030001 | metal ion transport | 1.61E-02 | 1.89E+00 | 1.29E-02 |
| GO:0006814 | GO:0006812 | monoatomic cation transport | 3.02E-02 | 1.41E+00 | 3.92E-02 |
| GO:0006814 | GO:0006811 | monoatomic ion transport | 2.30E-06 | 1.29E+00 | 5.19E-02 |
| GO:0006814 | GO:0055085 | transmembrane transport | 2.28E-02 | 1.12E+00 | 7.62E-02 |
| GO:0000413 | GO:0000413 | protein peptidyl-prolyl isomerization | 5.80E-05 | 3.31E+00 | 4.94E-04 |
| GO:0045089 | GO:0045089 | positive regulation of innate immune response | 6.70E-04 | 3.83E+00 | 1.47E-04 |
| GO:0045089 | GO:0002253 | activation of immune response | 2.56E-03 | 3.67E+00 | 2.15E-04 |
| GO:0045089 | GO:0009968 | negative regulation of signal transduction | 5.36E-03 | 2.94E+00 | 1.15E-03 |
| GO:0045089 | GO:0009967 | positive regulation of signal transduction | 3.93E-02 | 3.22E+00 | 6.08E-04 |
| GO:0010927 | GO:0010927 | cellular component assembly involved in morphogenesis | 6.90E-04 | 4.59E+00 | 2.58E-05 |
| GO:0010927 | GO:0022618 | protein-RNA complex assembly | 2.00E-02 | 2.69E+00 | 2.06E-03 |
| GO:0010927 | GO:0120031 | plasma membrane bounded cell projection assembly | 3.44E-03 | 3.58E+00 | 2.62E-04 |
| GO:0010927 | GO:0016358 | dendrite development | 2.56E-03 | 5.13E+00 | 7.50E-06 |
| GO:0010927 | GO:0007411 | axon guidance | 2.79E-03 | 3.73E+00 | 1.85E-04 |
| GO:0010927 | GO:0030041 | actin filament polymerization | 1.95E-03 | 4.40E+00 | 3.98E-05 |
| GO:0010927 | GO:0097435 | supramolecular fiber organization | 8.73E-03 | 3.13E+00 | 7.49E-04 |
| GO:0010927 | GO:0001745 | compound eye morphogenesis | 4.84E-03 | 5.43E+00 | 3.71E-06 |
| GO:0010927 | GO:0002009 | morphogenesis of an epithelium | 2.00E-02 | 3.89E+00 | 1.30E-04 |
| GO:0010927 | GO:0048707 | instar larval or pupal morphogenesis | 1.61E-02 | 8.47E+00 | 3.36E-09 |
| GO:0010927 | GO:0007051 | spindle organization | 9.01E-03 | 3.85E+00 | 1.42E-04 |
| GO:0010927 | GO:0061982 | meiosis I cell cycle process | 3.64E-02 | 4.10E+00 | 7.91E-05 |
| GO:0010927 | GO:0046530 | photoreceptor cell differentiation | 1.61E-02 | 5.17E+00 | 6.71E-06 |
| GO:0010927 | GO:0055001 | muscle cell development | 2.83E-02 | 4.58E+00 | 2.64E-05 |
| GO:0007610 | GO:0007610 | behavior | 1.15E-03 | 3.66E+00 | 2.18E-04 |
| GO:0007610 | GO:0009790 | embryo development | 3.84E-02 | 3.94E+00 | 1.14E-04 |
| GO:0007610 | GO:0032504 | multicellular organism reproduction | 3.68E-02 | 6.03E+00 | 9.41E-07 |
| GO:0007610 | GO:0002165 | instar larval or pupal development | 3.42E-03 | 5.80E+00 | 1.59E-06 |
| GO:0007610 | GO:0007552 | metamorphosis | 2.55E-02 | 5.91E+00 | 1.24E-06 |
| GO:0007424 | GO:0007424 | open tracheal system development | 1.33E-03 | 5.40E+00 | 4.00E-06 |
| GO:0007424 | GO:0009888 | tissue development | 1.80E-02 | 3.46E+00 | 3.50E-04 |
| GO:0007424 | GO:0048569 | post-embryonic animal organ development | 1.33E-02 | 5.13E+00 | 7.34E-06 |
| GO:0007424 | GO:0035220 | wing disc development | 1.61E-02 | 5.58E+00 | 2.65E-06 |
| GO:0007424 | GO:0040003 | chitin-based cuticle development | 5.65E-03 | 5.42E+00 | 3.85E-06 |
| GO:0007424 | GO:0061061 | muscle structure development | 3.14E-02 | 4.14E+00 | 7.34E-05 |
| GO:0006313 | GO:0006313 | DNA transposition | 3.67E-03 | 2.78E+00 | 1.66E-03 |
| GO:0006313 | GO:0006259 | DNA metabolic process | 4.01E-03 | 1.24E+00 | 5.81E-02 |
| GO:0007623 | GO:0007623 | circadian rhythm | 4.51E-03 | 4.02E+00 | 9.55E-05 |
| GO:1901615 | GO:1901615 | organic hydroxy compound metabolic process | 4.77E-03 | 2.16E+00 | 6.95E-03 |
| GO:0034655 | GO:0034655 | nucleobase-containing compound catabolic process | 5.59E-03 | 2.39E+00 | 4.11E-03 |
| GO:0034655 | GO:0046434 | organophosphate catabolic process | 3.95E-02 | 2.76E+00 | 1.75E-03 |
| GO:0060560 | GO:0060560 | developmental growth involved in morphogenesis | 8.31E-03 | 4.34E+00 | 4.58E-05 |
| GO:0060560 | GO:0040007 | growth | 2.82E-02 | 3.87E+00 | 1.36E-04 |
| GO:0008284 | GO:0008284 | positive regulation of cell population proliferation | 1.33E-02 | 3.59E+00 | 2.59E-04 |
| GO:0008284 | GO:0045944 | positive regulation of transcription by RNA polymerase II | 4.78E-02 | 3.15E+00 | 7.06E-04 |
| GO:0008284 | GO:0010628 | positive regulation of gene expression | 2.86E-02 | 2.81E+00 | 1.55E-03 |
| GO:0008284 | GO:0010638 | positive regulation of organelle organization | 8.31E-03 | 3.83E+00 | 1.50E-04 |
| GO:0008284 | GO:0051493 | regulation of cytoskeleton organization | 2.17E-02 | 3.26E+00 | 5.44E-04 |
| GO:0008284 | GO:1902903 | regulation of supramolecular fiber organization | 1.61E-02 | 3.41E+00 | 3.91E-04 |
| GO:0008284 | GO:0051128 | regulation of cellular component organization | 3.59E-02 | 2.53E+00 | 2.96E-03 |
| GO:0008284 | GO:0043254 | regulation of protein-containing complex assembly | 3.78E-02 | 3.42E+00 | 3.84E-04 |
| GO:0016051 | GO:0016051 | carbohydrate biosynthetic process | 2.46E-02 | 2.11E+00 | 7.74E-03 |
| GO:0051960 | GO:0051960 | regulation of nervous system development | 2.84E-02 | 4.00E+00 | 9.93E-05 |
| GO:0051960 | GO:0060284 | regulation of cell development | 2.84E-02 | 3.72E+00 | 1.89E-04 |
| GO:0051960 | GO:0022603 | regulation of anatomical structure morphogenesis | 3.95E-02 | 2.35E+00 | 4.43E-03 |
| GO:0055082 | GO:0055082 | intracellular chemical homeostasis | 2.87E-02 | 2.89E+00 | 1.28E-03 |
| GO:0007163 | GO:0007163 | establishment or maintenance of cell polarity | 3.78E-02 | 3.97E+00 | 1.06E-04 |

**Supplementary Table 5: Enriched GO terms for gene births at the Metamorphosis LCAs.**

| Cluster representative | Cluster member | Cluster member description | Member P-value | Member IC | Member frequency |
|---|---|---|---|---|---|
| GO:0019432 | GO:0019432 | triglyceride biosynthetic process | 3.50E-05 | 4.07E+00 | 8.51E-05 |
| GO:0050909 | GO:0050909 | sensory perception of taste | 1.00E-04 | 4.08E+00 | 8.34E-05 |
| GO:0043632 | GO:0043632 | modification-dependent macromolecule catabolic process | 2.10E-03 | 2.75E+00 | 1.76E-03 |
| GO:0043632 | GO:0009056 | catabolic process | 3.30E-02 | 1.50E+00 | 3.20E-02 |
| GO:0030036 | GO:0030036 | actin cytoskeleton organization | 2.20E-03 | 3.46E+00 | 3.49E-04 |
| GO:0030036 | GO:0044782 | cilium organization | 4.70E-02 | 3.59E+00 | 2.57E-04 |
| GO:0030036 | GO:0070286 | axonemal dynein complex assembly | 4.80E-02 | 4.27E+00 | 5.41E-05 |
| GO:0006399 | GO:0006399 | tRNA metabolic process | 3.20E-03 | 1.55E+00 | 2.80E-02 |
| GO:0006399 | GO:0006400 | tRNA modification | 2.64E-02 | 2.15E+00 | 7.11E-03 |
| GO:0006399 | GO:0000460 | maturation of 5.8S rRNA | 2.01E-02 | 4.46E+00 | 3.48E-05 |
| GO:0009308 | GO:0009308 | amine metabolic process | 3.30E-03 | 2.19E+00 | 6.41E-03 |
| GO:0009308 | GO:0009150 | purine ribonucleotide metabolic process | 2.11E-02 | 1.76E+00 | 1.73E-02 |
| GO:0071277 | GO:0071277 | cellular response to calcium ion | 8.60E-03 | 4.76E+00 | 1.76E-05 |
| GO:0031047 | GO:0031047 | RNA-mediated gene silencing | 1.77E-02 | 3.21E+00 | 6.19E-04 |
| GO:0031047 | GO:0043069 | negative regulation of programmed cell death | 2.64E-02 | 3.53E+00 | 2.96E-04 |
| GO:0031047 | GO:0045732 | positive regulation of protein catabolic process | 2.01E-02 | 3.69E+00 | 2.03E-04 |
| GO:0031047 | GO:0010608 | post-transcriptional regulation of gene expression | 3.33E-02 | 2.49E+00 | 3.25E-03 |
| GO:0060294 | GO:0060294 | cilium movement involved in cell motility | 2.17E-02 | 4.07E+00 | 8.60E-05 |
| GO:0006082 | GO:0006082 | organic acid metabolic process | 2.62E-02 | 9.77E-01 | 1.06E-01 |
| GO:0018401 | GO:0018401 | peptidyl-proline hydroxylation to 4-hydroxy-L-proline | 2.64E-02 | 5.35E+00 | 4.42E-06 |
| GO:0017157 | GO:0017157 | regulation of exocytosis | 3.36E-02 | 4.48E+00 | 3.32E-05 |
| GO:0060285 | GO:0060285 | cilium-dependent cell motility | 4.73E-02 | 4.55E+00 | 2.81E-05 |

**Supplementary Table 6: Enriched GO terms for gene births at the Sister LCAs.**

| Cluster representative | Cluster member | Cluster member description | Member P-value | Member IC | Member frequency |
|---|---|---|---|---|---|
| GO:0042593 | GO:0042593 | glucose homeostasis | 5.40E-04 | 4.00E+00 | 1.01E-04 |
| GO:0007417 | GO:0007417 | central nervous system development | 6.90E-04 | 4.63E+00 | 2.34E-05 |
| GO:0007417 | GO:0007422 | peripheral nervous system development | 9.50E-04 | 5.25E+00 | 5.65E-06 |
| GO:0007417 | GO:0035218 | leg disc development | 1.23E-02 | 6.39E+00 | 4.10E-07 |
| GO:0006952 | GO:0006952 | defense response | 8.50E-04 | 2.44E+00 | 3.64E-03 |
| GO:0006952 | GO:0070887 | cellular response to chemical stimulus | 5.10E-03 | 3.12E+00 | 7.62E-04 |
| GO:0099024 | GO:0099024 | plasma membrane invagination | 8.90E-04 | 5.06E+00 | 8.72E-06 |
| GO:0099024 | GO:0061024 | membrane organization | 3.42E-03 | 2.62E+00 | 2.40E-03 |
| GO:0030036 | GO:0030036 | actin cytoskeleton organization | 2.41E-03 | 3.46E+00 | 3.49E-04 |
| GO:0030036 | GO:0060271 | cilium assembly | 6.27E-03 | 3.61E+00 | 2.43E-04 |
| GO:0030036 | GO:0007015 | actin filament organization | 1.12E-02 | 3.21E+00 | 6.18E-04 |
| GO:0030036 | GO:0016050 | vesicle organization | 1.26E-02 | 3.79E+00 | 1.63E-04 |
| GO:0007416 | GO:0007416 | synapse assembly | 2.46E-03 | 4.58E+00 | 2.63E-05 |
| GO:0007416 | GO:0120031 | plasma membrane bounded cell projection assembly | 8.86E-03 | 3.58E+00 | 2.62E-04 |
| GO:0006006 | GO:0006006 | glucose metabolic process | 2.84E-03 | 2.50E+00 | 3.17E-03 |
| GO:0007156 | GO:0007156 | homophilic cell adhesion via plasma membrane adhesion molecules | 3.96E-03 | 3.36E+00 | 4.40E-04 |
| GO:0007156 | GO:0007155 | cell adhesion | 7.29E-03 | 2.55E+00 | 2.81E-03 |
| GO:0035290 | GO:0035290 | trunk segmentation | 4.35E-03 | 7.57E+00 | 2.69E-08 |
| GO:0035290 | GO:0016318 | ommatidial rotation | 5.00E-03 | 5.96E+00 | 1.10E-06 |
| GO:0010001 | GO:0010001 | glial cell differentiation | 4.59E-03 | 4.61E+00 | 2.46E-05 |
| GO:0010001 | GO:0007286 | spermatid development | 7.87E-03 | 4.75E+00 | 1.79E-05 |
| GO:0010001 | GO:0021700 | developmental maturation | 1.23E-02 | 4.25E+00 | 5.65E-05 |
| GO:0045595 | GO:0045595 | regulation of cell differentiation | 4.64E-03 | 3.35E+00 | 4.47E-04 |
| GO:0045595 | GO:0045572 | positive regulation of imaginal disc growth | 9.89E-03 | 6.01E+00 | 9.67E-07 |
| GO:0010035 | GO:0010035 | response to inorganic substance | 5.81E-03 | 3.26E+00 | 5.51E-04 |
| GO:0010035 | GO:1902074 | response to salt | 1.18E-02 | 3.86E+00 | 1.38E-04 |
| GO:0007218 | GO:0007218 | neuropeptide signaling pathway | 5.93E-03 | 3.74E+00 | 1.81E-04 |
| GO:0007218 | GO:0007254 | JNK cascade | 1.17E-02 | 4.94E+00 | 1.15E-05 |
| GO:0040003 | GO:0040003 | chitin-based cuticle development | 6.37E-03 | 5.42E+00 | 3.85E-06 |
| GO:0040003 | GO:0061061 | muscle structure development | 4.12E-03 | 4.14E+00 | 7.34E-05 |
| GO:0040003 | GO:0048568 | embryonic organ development | 5.13E-03 | 4.44E+00 | 3.62E-05 |
| GO:0032506 | GO:0032506 | cytokinetic process | 9.24E-03 | 2.94E+00 | 1.15E-03 |
| GO:0043409 | GO:0043409 | negative regulation of MAPK cascade | 9.25E-03 | 4.41E+00 | 3.91E-05 |
| GO:0043409 | GO:0010629 | negative regulation of gene expression | 1.32E-02 | 2.30E+00 | 5.04E-03 |
| GO:0007446 | GO:0007446 | imaginal disc growth | 1.03E-02 | 6.46E+00 | 3.46E-07 |
| GO:0071702 | GO:0055085 | transmembrane transport | 1.05E-02 | 1.12E+00 | 7.62E-02 |
| GO:0071702 | GO:0071702 | organic substance transport | 1.11E-02 | 1.45E+00 | 3.54E-02 |
| GO:0035151 | GO:0035151 | regulation of tube size, open tracheal system | 1.15E-02 | 5.50E+00 | 3.17E-06 |
| GO:0010506 | GO:0010506 | regulation of autophagy | 1.38E-02 | 3.68E+00 | 2.10E-04 |
| GO:0009267 | GO:0007268 | chemical synaptic transmission | 3.20E-05 | 3.83E+00 | 1.47E-04 |
| GO:0009267 | GO:0009267 | cellular response to starvation | 1.22E-02 | 3.46E+00 | 3.50E-04 |
| GO:0050877 | GO:0001700 | embryonic development via the syncytial blastoderm | 2.80E-04 | 5.82E+00 | 1.51E-06 |
| GO:0050877 | GO:0050877 | nervous system process | 1.04E-02 | 2.98E+00 | 1.06E-03 |
| GO:0008343 | GO:0008344 | adult locomotory behavior | 7.40E-04 | 4.76E+00 | 1.75E-05 |
| GO:0008343 | GO:0008343 | adult feeding behavior | 1.30E-02 | 5.14E+00 | 7.19E-06 |
| GO:0008343 | GO:0007619 | courtship behavior | 4.71E-03 | 5.39E+00 | 4.05E-06 |
| GO:0002009 | GO:0008586 | imaginal disc-derived wing vein morphogenesis | 3.87E-03 | 5.67E+00 | 2.13E-06 |
| GO:0002009 | GO:0002009 | morphogenesis of an epithelium | 1.16E-02 | 3.89E+00 | 1.30E-04 |
| GO:0032101 | GO:0032101 | regulation of response to external stimulus | 4.45E-03 | 2.89E+00 | 1.29E-03 |
| GO:0032101 | GO:0061057 | peptidoglycan recognition protein signaling pathway | 4.46E-03 | 6.03E+00 | 9.24E-07 |
| GO:0032101 | GO:0023056 | positive regulation of signaling | 5.20E-03 | 3.18E+00 | 6.57E-04 |
| GO:0016331 | GO:0008258 | head involution | 5.51E-03 | 5.85E+00 | 1.42E-06 |
| GO:0016331 | GO:0007391 | dorsal closure | 4.67E-03 | 5.41E+00 | 3.85E-06 |
| GO:0016331 | GO:0016331 | morphogenesis of embryonic epithelium | 7.41E-03 | 4.98E+00 | 1.06E-05 |
| GO:0016331 | GO:0007377 | germ-band extension | 1.30E-02 | 6.30E+00 | 5.01E-07 |
| GO:0008284 | GO:0051781 | positive regulation of cell division | 2.80E-04 | 4.12E+00 | 7.60E-05 |
| GO:0008284 | GO:0010647 | positive regulation of cell communication | 1.27E-02 | 3.18E+00 | 6.57E-04 |
| GO:0008284 | GO:0008284 | positive regulation of cell population proliferation | 6.84E-03 | 3.59E+00 | 2.59E-04 |
| GO:0008407 | GO:0007435 | salivary gland morphogenesis | 7.79E-03 | 5.37E+00 | 4.29E-06 |
| GO:0008407 | GO:0008407 | chaeta morphogenesis | 1.09E-02 | 5.78E+00 | 1.66E-06 |

**Supplementary Table 7: Semantic similarity clustering of enriched GO terms shared across gene families expanding at the Metamorphosis LCAs.**

| GO term | P-value | Description |
|---|---|---|
| GO:0006508 | 3.10E-06 | proteolysis |
| GO:0055085 | 0.00022 | transmembrane transport |
| GO:0019953 | 0.00033 | sexual reproduction |
| GO:0006869 | 0.00057 | lipid transport |
| GO:0071702 | 0.00251 | organic substance transport |
| GO:0007275 | 0.00669 | multicellular organism development |
| GO:0006836 | 0.01596 | neurotransmitter transport |
| GO:0044419 | 0.01844 | biological process involved in interspecies interaction between organisms |
| GO:0046483 | 0.02295 | heterocycle metabolic process |
| GO:1901565 | 0.03294 | organonitrogen compound catabolic process |
| GO:0031325 | 0.04065 | positive regulation of cellular metabolic process |
| GO:0050804 | 0.04447 | modulation of chemical synaptic transmission |

**Supplementary Table 8: Enriched GO terms shared across gene families expanding at the Sister LCAs.**

| GO term | P-value | Description |
|---|---|---|
| GO:0032501 | 0.00054 | multicellular organismal process |
| GO:0010629 | 0.00191 | negative regulation of gene expression |
| GO:0055085 | 0.00371 | transmembrane transport |
| GO:0071702 | 0.0068 | organic substance transport |
| GO:0030154 | 0.0166 | cell differentiation |
| GO:0070887 | 0.01952 | cellular response to chemical stimulus |

**Supplementary Table 9: Enriched GO terms shared across gene families expanding at the Deep nodes.**

| GO term | P-value | Description |
|---|---|---|
| GO:0015718 | 9.80E-08 | monocarboxylic acid transport |
| GO:0055085 | 1.60E-05 | transmembrane transport |
| GO:0006508 | 0.00052 | proteolysis |
| GO:0071702 | 0.001 | organic substance transport |
| GO:0070887 | 0.00239 | cellular response to chemical stimulus |
| GO:0071705 | 0.01081 | nitrogen compound transport |
| GO:0055082 | 0.01533 | intracellular chemical homeostasis |
| GO:0007015 | 0.01892 | actin filament organization |
| GO:0046394 | 0.02237 | carboxylic acid biosynthetic process |
| GO:0006796 | 0.0242 | phosphate-containing compound metabolic process |
| GO:0005975 | 0.02711 | carbohydrate metabolic process |
| GO:0006082 | 0.02738 | organic acid metabolic process |
| GO:0030855 | 0.02976 | epithelial cell differentiation |
| GO:0031032 | 0.03141 | actomyosin structure organization |
| GO:0006874 | 0.03807 | intracellular calcium ion homeostasis |
| GO:0042221 | 0.04257 | response to chemical |
| GO:0032501 | 0.044 | multicellular organismal process |
| GO:0006811 | 0.04741 | monoatomic ion transport |

**Supplementary Table 10: Enriched GO terms shared across gene families contracting at the Metamorphosis LCAs.**

| GO term | P-value | Description |
|---|---|---|
| GO:0006508 | 0.0034 | proteolysis |
| GO:0048856 | 0.0147 | anatomical structure development |
| GO:0030317 | 0.0156 | flagellated sperm motility |
| GO:0055085 | 0.0205 | transmembrane transport |
| GO:0006952 | 0.0225 | defense response |
| GO:0030308 | 0.047 | negative regulation of cell growth |

**Supplementary Table 11: Enriched GO terms shared across gene families contracting at the Sister LCAs.**

| Cluster representative | Cluster member | Cluster member description | Member P-value | Member IC | Member frequency |
|---|---|---|---|---|---|
| GO:0042593 | GO:0042593 | glucose homeostasis | 5.40E-04 | 4.00E+00 | 1.01E-04 |
| GO:0008586 | GO:0008586 | imaginal disc-derived wing vein morphogenesis | 3.87E-03 | 5.67E+00 | 2.13E-06 |
| GO:0008586 | GO:0007435 | salivary gland morphogenesis | 7.79E-03 | 5.37E+00 | 4.29E-06 |
| GO:0007156 | GO:0007156 | homophilic cell adhesion via plasma membrane adhesion molecules | 3.96E-03 | 3.36E+00 | 4.40E-04 |
| GO:0007156 | GO:0007155 | cell adhesion | 7.29E-03 | 2.55E+00 | 2.81E-03 |
| GO:0007391 | GO:0007391 | dorsal closure | 4.67E-03 | 5.41E+00 | 3.85E-06 |
| GO:0007391 | GO:0008258 | head involution | 5.51E-03 | 5.85E+00 | 1.42E-06 |
| GO:0007391 | GO:0008407 | chaeta morphogenesis | 1.09E-02 | 5.78E+00 | 1.66E-06 |
| GO:0008284 | GO:0008284 | positive regulation of cell population proliferation | 6.84E-03 | 3.59E+00 | 2.59E-04 |
| GO:0008284 | GO:0010629 | negative regulation of gene expression | 1.32E-02 | 2.30E+00 | 5.04E-03 |
| GO:0043409 | GO:0043409 | negative regulation of MAPK cascade | 9.25E-03 | 4.41E+00 | 3.91E-05 |
| GO:0055085 | GO:0055085 | transmembrane transport | 1.05E-02 | 1.12E+00 | 7.62E-02 |
| GO:0009267 | GO:0009267 | cellular response to starvation | 1.22E-02 | 3.46E+00 | 3.50E-04 |
| GO:0016318 | GO:0008344 | adult locomotory behavior | 7.40E-04 | 4.76E+00 | 1.75E-05 |
| GO:0016318 | GO:0016318 | ommatidial rotation | 5.00E-03 | 5.96E+00 | 1.10E-06 |
| GO:0016318 | GO:0035290 | trunk segmentation | 4.35E-03 | 7.57E+00 | 2.69E-08 |
| GO:0007416 | GO:0061024 | membrane organization | 3.42E-03 | 2.62E+00 | 2.40E-03 |
| GO:0007416 | GO:0007416 | synapse assembly | 2.46E-03 | 4.58E+00 | 2.63E-05 |
| GO:0007416 | GO:0060271 | cilium assembly | 6.27E-03 | 3.61E+00 | 2.43E-04 |
| GO:0040003 | GO:0007417 | central nervous system development | 6.90E-04 | 4.63E+00 | 2.34E-05 |
| GO:0040003 | GO:0007422 | peripheral nervous system development | 9.50E-04 | 5.25E+00 | 5.65E-06 |
| GO:0040003 | GO:0040003 | chitin-based cuticle development | 6.37E-03 | 5.42E+00 | 3.85E-06 |

**Supplementary Table 12: Semantic similarity clustering of enriched GO terms shared across gene families expanding at the Metamorphosis LCAs and showing signatures of adaptive lineage-specific expansion.**

| Family label | GO term | Drosophila melanogaster protein identifier |
|---|---|---|
| Family 1 | GO:0002143,GO:0005515,GO:0005829,GO:0005886,GO:0007156,GO:0008046, GO:0008641,GO:0016783,GO:0030424,GO:0043025,GO:0050808,GO:0070566, GO:0005911,GO:0050839,GO:0098609,GO:0043005,GO:0019897,GO:0045466, GO:0007465,GO:0048149,GO:0007615,GO:0007616,GO:0003676,GO:0016787, GO:0046872,GO:0003993,GO:0007155,GO:0007411,GO:0070593,GO:0098632, GO:0006801,GO:0032589 | NP_524454.2,NP_001287571.1,NP_649339.2,NP_611666.2 |
| Family 2 | GO:0000226,GO:0000278,GO:0003924,GO:0005200,GO:0005525,GO:0005737, GO:0005874,GO:0007017,GO:0007435 | NP_524290.2,NP_725896.1 |
| Family 3 | GO:0016477,GO:0007155,GO:0035147,GO:0008045,GO:0007436,GO:0005515, GO:0005886,GO:0005615,GO:0031012,GO:0007166,GO:0007411,GO:0007156, GO:0072499,GO:0036194,GO:0007416,GO:0035230 | NP_001261796.1,NP_001303391.1 |
| Family 4 | GO:0016020,GO:0022857,GO:0055085 | NP_650818.1,NP_650391.1,NP_650819.1,NP_650815.2,NP_650814.2, NP_650390.2,NP_647998.2,NP_650817.1,NP_650392.1,NP_647996.1,NP_648019.2, NP_650816.3 |
| Family 5 | GO:0008345,GO:0008344,GO:0007540,GO:0007549,GO:0005634,GO:0000122, GO:0001227,GO:0045840,GO:0006355,GO:0003677,GO:0046983,GO:0002052, GO:0014019,GO:1902692,GO:0007465,GO:0000987,GO:0043565,GO:1990837, GO:0003700,GO:0010629,GO:0005829,GO:0000978,GO:0050767,GO:0000981, GO:0006357,GO:0009952,GO:0045892,GO:0070888,GO:0005515,GO:0001222, GO:0007366,GO:0000902,GO:0035239,GO:0007435,GO:0061024,GO:0007399, GO:0035290,GO:0035289,GO:0007424,GO:0001666 | NP_476923.1,NP_523977.2 |
| Family 6 | GO:0003924,GO:0005525,GO:0005886,GO:0007165,GO:0019003,GO:0007265, GO:0007173,GO:0005634,GO:0000785,GO:0045815,GO:0070828,GO:0003682, GO:0000812,GO:0031491,GO:0005575,GO:0008150,GO:0003674,GO:0006357, GO:0003700,GO:0000978,GO:0050905,GO:0051607,GO:0005515,GO:0016020, GO:0007362,GO:0008293,GO:0007465,GO:0045500,GO:0007427,GO:0000165, GO:0042461,GO:0043703,GO:0008543,GO:0001710,GO:2001234,GO:0045465, GO:0048626,GO:0007298,GO:0046530,GO:0048749,GO:0030381,GO:0007474, GO:0070374,GO:0048010,GO:0035099,GO:0046664,GO:0007472,GO:0007476, GO:0007422,GO:0007369,GO:0016318,GO:0040008,GO:0046673,GO:0007479, GO:0002168,GO:0040014,GO:0007426,GO:0019887,GO:0043539,GO:0003925, GO:0007455,GO:0043066,GO:0035088,GO:0007309,GO:0008284,GO:0046534, GO:0016242,GO:0009267,GO:0008361,GO:0007552,GO:0045793,GO:0048865, GO:0048863,GO:0072089,GO:0072002,GO:0008594,GO:0001709,GO:0008586, GO:0042981,GO:0060438,GO:0036335,GO:1904263,GO:0035169,GO:0035170, GO:0007430,GO:0008340,GO:0007395,GO:0010629,GO:0007391,GO:0035208, GO:0035313 | NP_523917.2,NP_476699.1 |
| Family 7 | GO:0000132,GO:0008093,GO:0008356,GO:0009786,GO:0045176,GO:0045179, GO:0005515,GO:0007525,GO:0007422,GO:0040001,GO:0008104,GO:0006403, GO:0005737,GO:0061382,GO:0005938,GO:0007423,GO:0055059,GO:0051960, GO:0050829, GO:0007400,GO:1990499 | NP_477024.1 |
| Family 8 | GO:0006629,GO:0016788,GO:0016042,GO:0004806,GO:0004620,GO:0047372, GO:0017171,GO:0005811,GO:0005615,GO:0019953,GO:0042593,GO:0033993, GO:0016298,GO:0005576,GO:0007586,GO:0005102,GO:0007275,GO:0016055 | NP_477331.1,NP_609428.1,NP_001260356.1,NP_652714.2,NP_611020.1, NP_650218.3,NP_523540.1,NP_001014548.1,NP_609420.1 |
| Family 9 | GO:0005515,GO:0003735,GO:0005840,GO:0006412,GO:0022625,GO:0005615, GO:0031012,GO:0005834,GO:0007200,GO:0031681,GO:0050909,GO:0005886, GO:0030514,GO:0000045,GO:0016236,GO:0035096,GO:0061723,GO:0009267, GO:0060786,GO:0035883,GO:0034274,GO:0090385,GO:0000423,GO:0061709, GO:0034045,GO:0043495,GO:0000421,GO:0000123,GO:0044545,GO:0005634, GO:0045944,GO:0061174,GO:0043121,GO:0030424,GO:0048167,GO:0005030, GO:0061630,GO:0097602,GO:0005680,GO:0008270,GO:0031145,GO:0045842, GO:0000151,GO:0016567,GO:0006511, GO:0008150,GO:0003674,GO:0005575 | NP_609615.2,NP_573382.1,NP_001263151.1 |
| Family 10 | GO:0004497,GO:0005506,GO:0016705,GO:0020037,GO:0007494,GO:0007391, GO:0008362,GO:0008258,GO:0006697,GO:0042767,GO:0007417,GO:0005739, GO:0005743,GO:0006700,GO:0006704,GO:0008203,GO:0034650,GO:0071375 , GO:0017085, GO:0007490,GO:0005737 | NP_524810.2,NP_611370.2,NP_650783.2,NP_650782.1,NP_610796.1, NP_610588.2,NP_649052.1 |
| Family 11 | GO:0003700,GO:0006355,GO:0008270,GO:0043565,GO:0000785,GO:0004879, GO:0005634,GO:0006357,GO:0034056,GO:0045892,GO:0007427,GO:0007088, GO:0007417,GO:0001227,GO:0140297,GO:0035290,GO:0007354,GO:0005515 | NP_788552.1,NP_524206.1,NP_001287130.1 |
| Family 12 | GO:0001664,GO:0002031,GO:0005737,GO:0007165,GO:0002046,GO:0016060, GO:0016028,GO:0016059,GO:0007608,GO:0002029,GO:0006897,GO:0045494, GO:0005515,GO:0016029,GO:0007605,GO:0060271,GO:0035869,GO:0036038, GO:0008150,GO:0003674,GO:0005575,GO:0036094,GO:0005813,GO:0007098, GO:0035148,GO:0031023,GO:0030010,GO:0005814,GO:0001764,GO:0005615, GO:0007186,GO:0045746,GO:0035641,GO:0048260,GO:0045751,GO:0120177, GO:0043409,GO:0005829,GO:0045879,GO:0031648,GO:0060189,GO:0050728, GO:0048149,GO:0035207,GO:0007616,GO:0007131,GO:0005524,GO:0003677, GO:0006270,GO:0005634,GO:0008094,GO:0003678,GO:0003697,GO:0017116, GO:0042555 | NP_476681.1,NP_523976.1,NP_524988.1 |
| Family 13 | GO:0005515,GO:0005615,GO:0007165,GO:0031012,GO:0009653,GO:0004888, GO:0005886,GO:0007155,GO:0019731,GO:0006955,GO:0019730,GO:0030703, GO:0007297,GO:0060026,GO:0003401,GO:0007419,GO:0005576,GO:0008200, GO:0060049,GO:0048935,GO:0048018,GO:0005030,GO:0070983,GO:0048167, GO:0008201,GO:0048495,GO:0002224,GO:0006954,GO:0002752,GO:0009897, GO:0051607,GO:0038187,GO:0046790,GO:0010508,GO:0050688,GO:0007411, GO:0008063,GO:0009597,GO:0002225,GO:0045087 | NP_476814.1,NP_524757.1,NP_524081.1,NP_523797.1 |
| Family 14 | GO:0005615,GO:0062023,GO:0007399,GO:0008407,GO:0048749,GO:0016318, GO:0008587,GO:0097305 | NP_476710.2 |
| Family 15 | GO:0005615,GO:0031012,GO:0008010,GO:0040003,GO:0062129,GO:0004252, GO:0006508,GO:0008233,GO:0045087,GO:0008286,GO:0005158,GO:0005634, GO:0006355,GO:0003700,GO:0046983,GO:0051726,GO:0043565,GO:0030308, GO:0000122,GO:0046982,GO:0005515,GO:0005654,GO:0000981,GO:0006357, GO:0000978 | NP_647668.1,NP_649678.1,NP_001137869.1,NP_648209.1,NP_001097547.1 |

**Supplementary Table 13: Gene families with signatures of adaptive lineage-specific gene family expansions at the Metamorphosis LCAs.**
